# A Mesoscale Framework for Psychedelic Drug Action in the Human Brain

**DOI:** 10.1101/2025.11.23.690016

**Authors:** Rui Dai, Rodrigo Cofré, Christopher Timmermann, Robin L. Carhart-Harris, Anthony G. Hudetz, Zirui Huang, George A. Mashour

## Abstract

The mechanism of psychedelic drug action is a dynamic area of neuroscience, with two major lines of investigation: (1) laboratory studies at the molecular and cellular level, and (2) human neuroimaging studies of functional brain networks. Despite considerable progress, there remains insufficient understanding of the link between molecular/cellular substrates of psychedelics and the whole-brain network effects that result. Here we report a study of psychedelic action that focuses on the intermediate spatial scale of local brain regions (<1cm^3^). We analyzed the effects of classical psychedelics (dimethyltryptamine [DMT], lysergic acid diethylamide [LSD], psilocybin) and non-classical psychedelics (nitrous oxide, ketamine) in humans using functional magnetic resonance imaging. We found that all five drugs reduced regional homogeneity, that is, they disrupted local synchrony, in small-scale brain regions; this disruption occurred extensively in cortical regions and sparsely in subcortical regions. Dynamic analysis of both regional homogeneity and global functional connectivity showed an inverse pattern, with large-scale functional connectivity being enhanced as local synchrony declined. We then conducted dominance analysis to assess the contribution of various neurotransmitter receptors to changes in regional homogeneity. DMT, LSD, and psilocybin showed the 5-HT receptors as the most dominant association; by contrast, regional homogeneity changes attributable to both nitrous oxide and ketamine were most strongly associated with the NMDA receptor. Both neuronal (including interneurons) and non-neuronal cell types were linked to psychedelic-induced changes in synchrony at the level of local brain regions. These data, across five drugs from two drug classes, provide evidence that a diverse set of molecular and cellular events lead to a common outcome of disrupted synchrony in local brain regions, which in turn mediate drug-specific changes in global functional connectivity.

## Introduction

Psychedelics are potent psychoactive compounds that produce profound alterations in perception, emotion, and cognition. Renewed scientific interest in psychedelics stems from promising evidence of therapeutic efficacy in psychiatric conditions where conventional treatments often fail. The spectrum of psychedelic agents includes the so-called “classical” psychedelic drugs, such as DMT, LSD, and psilocybin, which are thought to work primarily through serotonergic 5-HT₂A receptors, and atypical psychedelics such as ketamine and nitrous oxide, with distinct molecular targets. These compounds have shown therapeutic potential for depression, bipolar disorder, anxiety, post-traumatic stress disorder, and addiction ^1–7^. Although the psychoactive effects of psychedelics have attracted widespread attention, their underlying neurobiological mechanisms remain incompletely understood. Identifying consistent neural signatures across diverse psychedelic compounds may provide crucial insights into fundamental mechanisms, despite their pharmacological diversity.

To date, two major lines of investigation have dominated the field. On the one hand, laboratory-based studies in animal models have characterized the molecular and cellular pathways engaged by psychedelics, identifying key receptors such as serotonin 5-HT₂A ^8,9^, NMDA ^10,11^ and others ^12,13^, and mapping their downstream signaling cascades ^8,14–17^. On the other hand, neuroimaging studies in humans have revealed large-scale changes in brain networks, including increased global functional connectivity and reorganization of canonical resting-state networks^18–22^. However, a critical gap remains in understanding how receptor- and cell-level mechanisms induced by psychedelics propagate to large-scale brain network reconfigurations observed in humans. We therefore focused this investigation on the *mesoscale*, which encompasses local circuit dynamics and intra-regional coordination, and which represents a crucial but underexplored level of organization. Few studies have systematically investigated how psychedelics modulate activity and integration at this intermediate scale, leaving a key link in the mechanistic bridge between molecular action and altered conscious experience unresolved.

Regional homogeneity (ReHo)^23^, a voxel-based resting-state functional magnetic resonance imaging (fMRI) metric of local neural synchrony, provides a mesoscale index ideally suited to bridge the gap between molecular mechanisms and large-scale functional network changes. ReHo quantifies the synchrony of low-frequency BOLD signal fluctuations between a given voxel and its neighboring voxels. For example, under sustained visual stimulation, ReHo in the visual cortex increases. Although ReHo and task activation can be correlated ^24^, they capture distinct aspects of neural dynamics ^23,25^. ReHo reflects changes in local temporal synchrony among neighboring voxels, whereas task activation represents changes in overall neural activity estimated using a hemodynamic response–based model fit ^23^. Thus, ReHo provides a task-free, data-driven assessment of regional functional organization without relying on a priori assumptions, making it particularly advantageous for uncovering spatially heterogeneous, compound-specific effects of drugs on the brain ^23,24^. However, its application in psychedelic research remains limited, with only a few studies examining ketamine^26–28^ and little to no work on serotonergic compounds or other atypical agents such as nitrous oxide. Moreover, it remains unclear whether local changes in ReHo are associated with specific receptors profiles or shaped by the distribution of distinct neuronal and non-neuronal cell types. Elucidating these relationships is essential for developing integrative models of psychedelic action that span molecular-, cellular-, and systems-level organization, and may ultimately inform the design of more targeted and effective therapeutic interventions. By leveraging ReHo as a bridge between molecular signaling and macroscale dynamics, the present study offers a novel entry point to dissect the mesoscale mechanisms of psychedelic action.

Here, we address this gap of knowledge by combining fMRI with receptor-enrichment and cell-type association analyses to examine the impact of both classical (DMT, LSD, psilocybin) and non-classical (ketamine and nitrous oxide) psychedelics on local brain synchrony in humans. We first tested whether all five drugs disrupt ReHo in cortical and subcortical areas and examined how these local effects dynamically co-occurred with changes in global functional connectivity. We then applied dominance analysis, a method that quantifies the relative importance of predictors in explaining a dependent variable, to assess the contribution of different neurotransmitter receptors to local synchrony changes and to examine the potential involvement of distinct neuronal and non-neuronal cell types. Lastly, we conducted a mediation analysis to assess whether changes in ReHo mediate the relationship between molecular-level perturbations and large-scale alterations in global functional connectivity.

## Results

### Psychedelics reduce local synchrony across cortical networks

We analyzed fMRI data from 63 healthy volunteers across five independent studies to quantify local neural synchrony using ReHo to quantify changes in local neural synchrony induced by the psychedelic compounds DMT (n = 14), LSD (n = 15), psilocybin (n = 7), ketamine (n = 12), and nitrous oxide (n = 15). ReHo captures the similarity of fMRI time series among neighboring voxels by computing Kendall’s coefficient of concordance (W) within a 27-voxel neighborhood, generating a voxel-wise measure of local coherence. For each subject, ReHo values were averaged globally and within canonical functional networks to quantify local coherence at multiple spatial levels (Figure 1).

**Figure 1:**
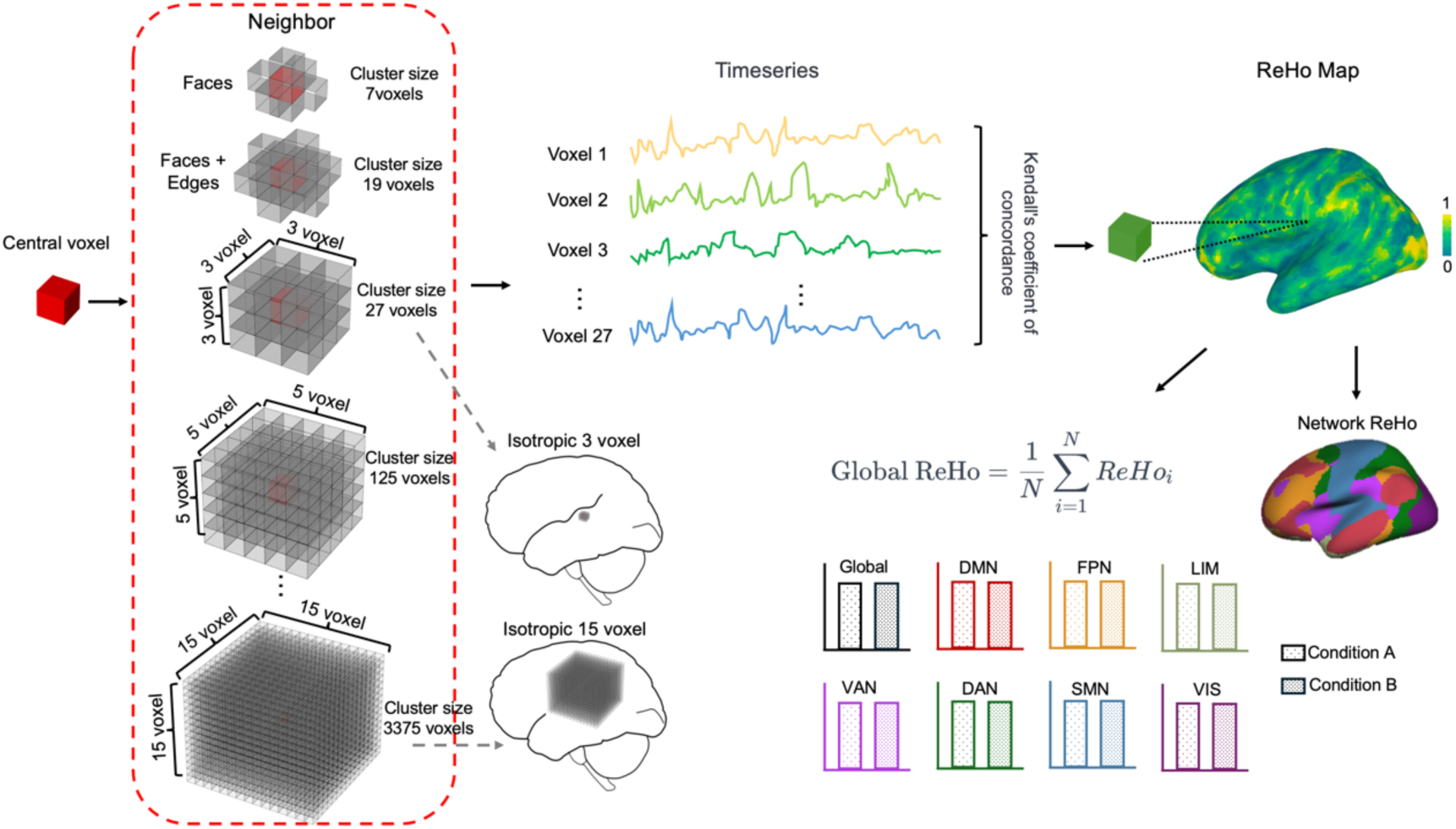
Schematic illustration of the methodology of regional homogeneity (ReHo) calculation, starting from the voxel level to the aggregation of global and network ReHo measures. ReHo measures concordance, a nonparametric metric of synchrony among adjacent voxel time series. For each voxel in the gray matter (center voxel in the figure), a Kendall’s correlation coefficient value is computed between that voxel’s timeseries and its 26 neighbors who share at least a face, edge, or corner with the center voxel. This voxel-wise calculation is then averaged to derive global and network-specific ReHo measures. Additionally, variations in ReHo cluster sizes in the analysis are depicted, as well as the portion of cluster in the brain. DMN: default-mode network, FPN: frontoparietal network, LIM: limbic network, VAN: ventral-attention network, DAN: dorsal-attention network, SMN: somatomotor network, VIS: visual network.

All five psychedelics reduced global average ReHo compared to baseline, representing an overall cortical desynchrony (Figure 2). At the network level, DMT showed significant ReHo reductions in DMN, FPN, VAN, DAN, VIS, and SMN, but not in LIM. LSD produced the most widespread effects, with significant reductions across all seven networks. Psilocybin produced moderate but consistent reductions, with the strongest effects in the FPN and DMN, followed by VIS, DAN, and VAN, while changes in LIM and SMN were small and not significant. ReHo reductions with ketamine and nitrous oxide were slightly smaller in magnitude than those observed with DMT, LSD, and psilocybin but remained widespread. Ketamine showed significant decreases in FPN, DAN, DMN, VAN, SMN, and VIS, while LIM showed minimal and nonsignificant change. Nitrous oxide exhibited the same pattern—significant reductions in FPN, DMN, DAN, VIS, SMN, and VAN, with LIM not significant (Table 1).

**Figure 2:**
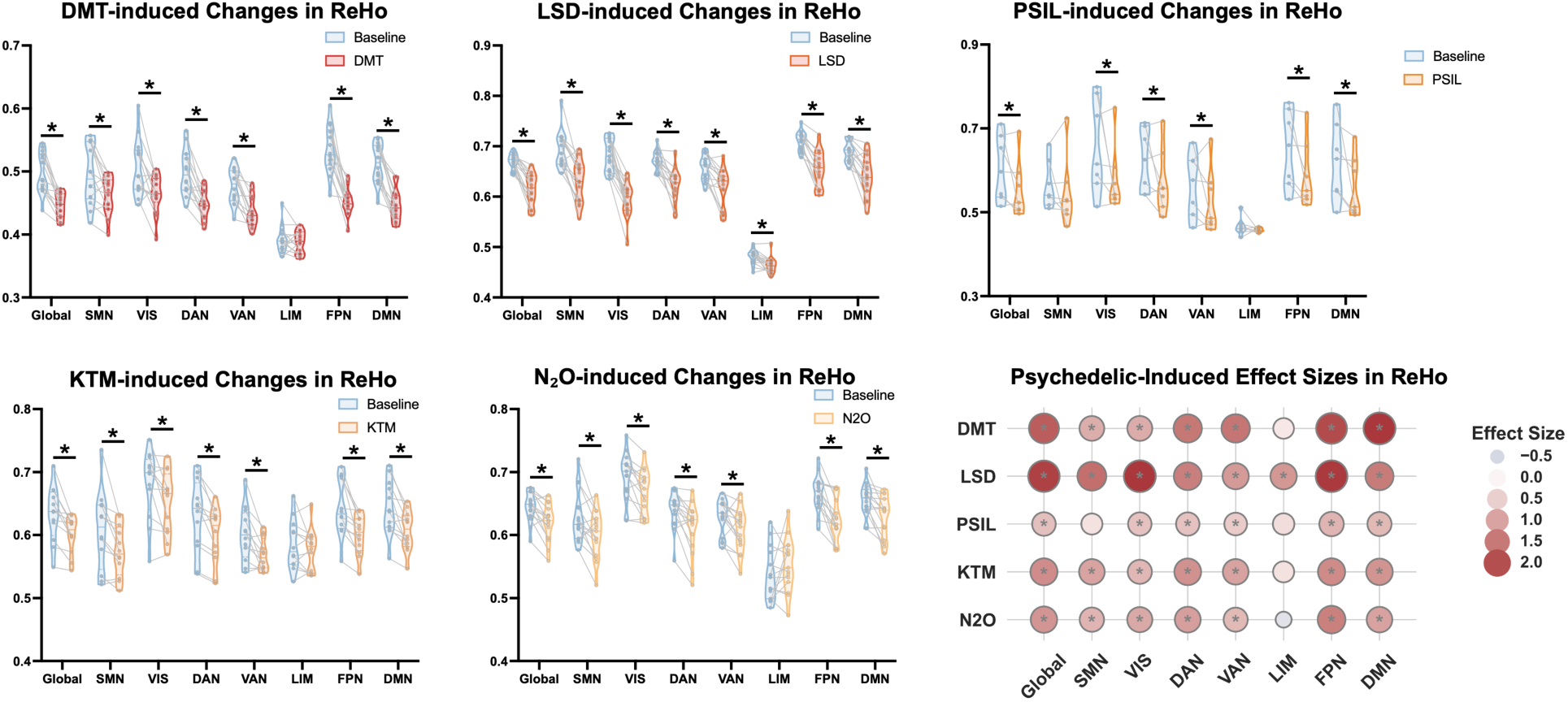
Regional homogeneity (ReHo) changes across cortical average and seven cortical networks in baseline versus acute psychedelic conditions. Violin plots (left) show global and network-specific ReHo values for each drug compared to baseline (SMN: somatomotor network; VIS: visual network; DAN: dorsal attention network; VAN: ventral attention network; LIM: limbic network; FPN: frontoparietal network; DMN: default-mode network). Bubble plots (right) display effect sizes (Cohen’s d) of ReHo changes (baseline vs. drug); bubble size and color represent effect magnitude. Asterisks denote significant ReHo reductions with FDR corrected p<0.05. DMT: dimethyltryptamine, LSD: lysergic acid diethylamide, PSIL: psilocybin, KTM: ketamine, N_2_O: nitrous oxide.

**Table 1.** Cohen’s d effect sizes and FDR-corrected p-values (pFDR) for ReHo reductions across cortical networks.

|  | DMT | LSD | PSIL | KTM | N<sub>2</sub>O |
| --- | --- | --- | --- | --- | --- |
| **GLOBAL** | d = 1.82, pFDR < 0.001 | d = 2.13, pFDR < 0.001 | d = 0.67, pFDR = 0.005 | d = 1.26, pFDR = 0.004 | d = 1.14, pFDR = 0.002 |
| **SMN** | d = 0.86, pFDR = 0.008 | d = 1.61, pFDR < 0.001 | d = 0.29, pFDR = 0.210 | d = 1.00, pFDR = 0.007 | d = 0.79, pFDR = 0.011 |
| **VIS** | d = 0.89, pFDR = 0.007 | d = 2.23, pFDR < 0.001 | d = 0.64, pFDR = 0.006 | d = 0.76, pFDR = 0.027 | d = 0.92, pFDR = 0.005 |
| **DAN** | d = 1.49, pFDR < 0.001 | d = 1.40, pFDR < 0.001 | d = 0.63, pFDR = 0.006 | d = 1.20, pFDR = 0.004 | d = 1.04, pFDR = 0.003 |
| **VAN** | d = 1.52, pFDR < 0.001 | d = 1.10, pFDR < 0.001 | d = 0.53, pFDR = 0.023 | d = 1.01, pFDR = 0.007 | d = 0.75, pFDR = 0.013 |
| **LIM** | d = 0.22, pFDR = 0.417 | d = 1.13, pFDR < 0.001 | d = 0.35, pFDR = 0.183 | d = 0.29, pFDR = 0.337 | d = -0.36, pFDR = 0.185 |
| **FPN** | d = 2.00, pFDR < 0.001 | d = 2.20, pFDR < 0.001 | d = 0.81, pFDR = 0.002 | d = 1.27, pFDR = 0.004 | d = 1.38, pFDR < 0.001 |
| **DMN** | d = 2.24, pFDR < 0.001 | d = 1.44, pFDR < 0.001 | d = 0.75, pFDR = 0.002 | d = 1.17, pFDR = 0.004 | d = 1.00, pFDR = 0.003 |

Thus, each psychedelic induced significant ReHo reductions across cortical networks, with the most consistent effects observed in high-order association networks such as the default mode (DMN) and frontoparietal (FPN) networks. Effects in the limbic network were less robust, which could reflect known limitations in signal quality within this region.

In subcortical regions, ReHo reductions were generally smaller and less consistent across drugs (Figure S1). After FDR correction, only LSD showed significant decreases, most prominently in the anterior thalamus, posterior thalamus, and hippocampus as well as in the subcortical average. Several additional subcortical regions (e.g., caudate, globus pallidus, and nucleus accumbens) showed minimal decreases that did not survive FDR correction. Psilocybin also induced significant reductions in the posterior thalamus, nucleus accumbens, and globus pallidus, with additional minimal effects in the anterior thalamus, amygdala, and subcortical average. DMT and N₂O exhibited minimal reductions in the caudate, whereas ketamine showed no significant subcortical effects.

### ReHo alterations are most prominent at fine spatial resolution

Because the spatial scale of local synchrony may influence how psychedelic effects manifest across cortical and subcortical regions, we systematically varied the neighborhood size for ReHo computation to evaluate the spatial-scale dependence of psychedelic-induced ReHo changes. Specifically, neighborhoods were defined as: 7 voxels (central voxel plus 6 face neighbors, ≈ 1-voxel radius, ∼3 mm), 19 voxels (central voxel plus 18 face/edge neighbors), and cubic neighborhoods of increasing size from 27 voxels (3×3×3 cube, radius = 1 voxel ≈ 3 mm) up to 3375 voxels (15×15×15 cube, radius = 7 voxels ≈ 21 mm). Across all five drugs, we observed that ReHo reductions were most pronounced at neighborhoods (7–27 voxels), with effect sizes diminishing as the spatial scale increased (Figure 3, Table S1- S5). This pattern was consistent in both global and network-level analyses.

**Figure 3:**
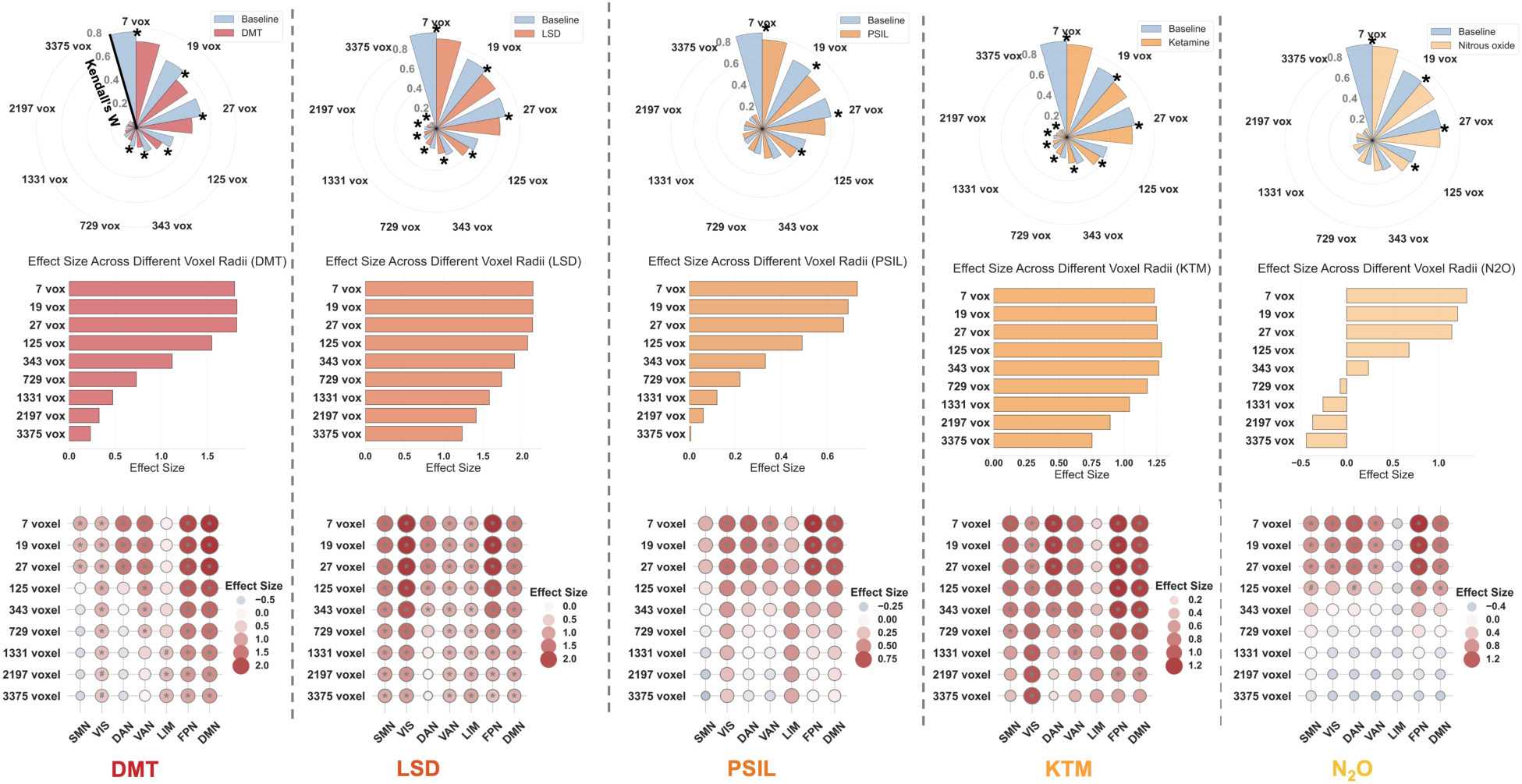
Spatial-scale dependence of psychedelic-induced regional homogeneity (ReHo) changes. For each drug (dimethyltryptamine/DMT, lysergic acid diethylamide/LSD, psilocybin/PSIL, ketamine/KTM, and nitrous oxide/N_2_O), ReHo was computed using neighborhoods of increasing size: 7 voxels (central voxel + 6 face neighbors), 19 voxels (central voxel + 18 face/edge neighbors), and cubic neighborhoods from 27 voxels (3×3×3 cube, radius = 1 voxel ≈ 3 mm) up to 3375 voxels (15×15×15 cube, radius = 7 voxels ≈ 21 mm). Polar plots (top) show global ReHo values for drug versus baseline across radii, with asterisks indicating significant reductions after FDR correction (p<0.05). Bar plots (middle) display effect sizes (Cohen’s d) of global ReHo changes across radii. Bubble plots (bottom) illustrate network-specific effect sizes across spatial scales (SMN: somatomotor network; VIS: visual network; DAN: dorsal attention network; VAN: ventral attention network; LIM: limbic network; FPN: frontoparietal network; DMN: default-mode network).

Global ReHo under DMT decreased most prominently at fine spatial scales, with significant reductions across the first six neighborhood sizes. LSD yielded the strongest and most pervasive global decreases, which were significant at all spatial scales. Psilocybin showed moderate global reductions, limited to the smallest neighborhood sizes. Ketamine produced consistent reductions across all scales, each surviving FDR correction. Nitrous oxide exhibited weaker global effects, with significant reductions only at the smallest four neighborhood sizes (Table S1- S5).

At the network level, scale-dependent ReHo reductions were evident across all psychedelics, with the strongest effects observed consistently in higher-order cortical networks such as the DMN and FPN, particularly at smaller voxel radii. For DMT, the largest decreases occurred in the DMN and FPN, with moderate reductions in DAN and VAN, and minimal changes in LIM. LSD produced the most widespread and robust reductions, with the strongest effects in VIS, FPN, and DMN, alongside substantial decreases in other cortical networks. Psilocybin induced more moderate and spatially constrained reductions, with significant effects at fine spatial scales in FPN and DMN, followed by DAN, VIS and VAN, while SMN and LIM showed weaker and non-significant changes. For ketamine, the largest reductions were again observed in the DMN and FPN, with smaller but consistent decreases across other networks. Nitrous oxide exhibited the weakest and most variable network-level changes, with notable reductions in FPN and DAN at fine spatial scales, and minimal effects in most other networks. Overall, these results demonstrate that psychedelic-induced ReHo reductions are most pronounced in association networks (DMN, FPN) at fine spatial resolutions, with effect sizes declining markedly as the cluster radius increases (Table 2-6). Subcortical ReHo reductions were most evident with LSD, particularly at smaller voxel radii, while the other compounds showed weaker or more variable subcortical effects (see Supplementary Figure 2).

Together, these results suggest that local desynchronization induced by psychedelics occurs predominantly within fine-grained, mesoscopic neural circuits (Figure S3). The data reinforce the interpretation of ReHo as a scale-sensitive measure and underscore the importance of spatial resolution when probing local changes under altered states of consciousness.

### Psychedelics induce dynamically dissociable changes in local and global connectivity

Although average ReHo showed robust reductions across the entire scan duration, these changes do not capture the moment-to-moment fluctuations in local neural synchrony that may be critical to understanding the psychedelic experience. To gain insight into dynamic changes in regional synchrony, we applied a sliding-window approach to compute dynamic ReHo (dReHo), enabling the assessment of how local synchrony evolves throughout the course of psychedelic exposure. In addition to characterizing the temporal dynamics of local synchrony itself, we also compared its relationship with global functional connectivity (GFC)—a commonly used marker of global brain integration that has been shown to increase with psychedelics ^19,21,22,29,30^. Prior studies have analyzed dynamic GFC (dGFC), but the temporal relationship between dReHo and dGFC has not been explored. Our analysis thus sought to assess whether local and global dynamics evolve in concert or in dissociation under the influence of different compounds.

Across all five psychedelics, we observed a consistent pattern of pronounced reductions in local synchrony and comparatively modest increases in global connectivity (Figure 4). During DMT administration, dReHo decreased sharply following drug onset and remained suppressed throughout the scan, while placebo showed only a brief decrease. In contrast, dGFC increased in both DMT and placebo conditions, and the between-condition difference in mean dGFC was not statistically significant after permutation correction. In the LSD condition, dReHo showed a sustained and statistically significant reduction compared to placebo, paralleling a persistent increase in dGFC. In the psilocybin condition, dReHo was markedly reduced relative to baseline, while dGFC showed a modest increase. However, due to the limited sample size (n = 7), the temporal clusters of significance did not survive FWE correction; uncorrected effects are shown in light gray shading. Ketamine produced a gradual decline in dReHo over the course of the drug phase, accompanied by a steady increase in dGFC. With nitrous oxide, dReHo exhibited a clear reduction, while dGFC rose gradually across time. However, permutation testing did not reveal a significant group difference in dGFC between N₂O and baseline conditions, indicating a less pronounced global effect for this compound. Together, these results suggest that psychedelics dynamically reorganize brain activity by dampening local coherence while enhancing large-scale functional integration.

**Figure 4:**
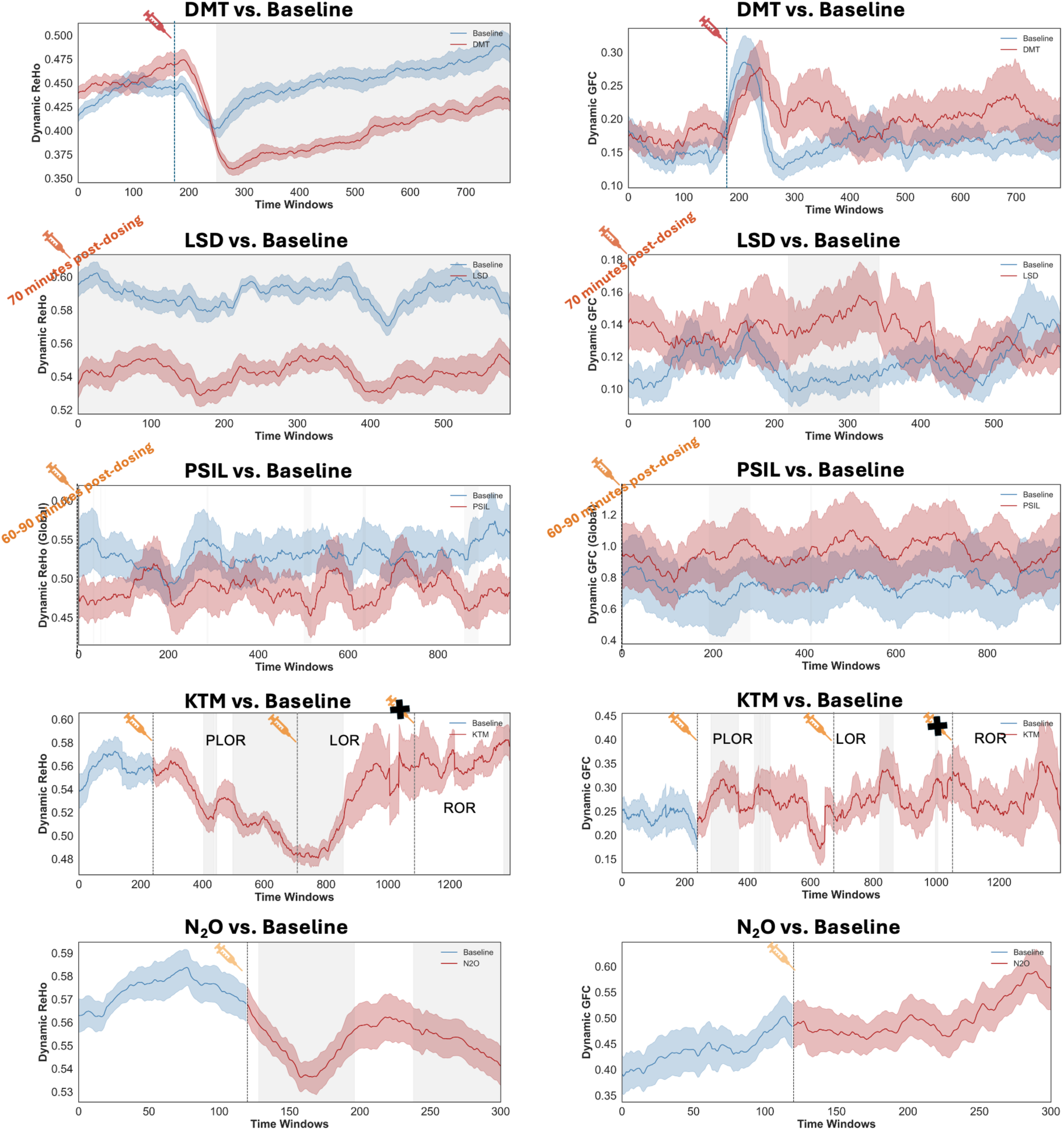
Dynamic regional homogeneity (dReHo) and global functional connectivity (GFC) pre- and post-exposure to psychedelics. dReHo (left) and dynamic GFC (right) were computed using a sliding-window approach (window = 60 TRs, step = 1 TR). For each drug (dimethyltryptamine/DMT, lysergic acid diethylamide/LSD, psilocybin/PSIL, ketamine/KTM, and nitrous oxide/N_2_O), solid lines represent group means, with shaded bands indicating the standard error. Statistical significance of drug–baseline differences was assessed at each time point using paired comparisons across subjects. Clusters of consecutive time windows exceeding the nominal threshold (p < 0.05, two-tailed) were identified, and cluster significance was determined via permutation testing with 5,000 random condition-label shuffles. Gray bars denote consecutive time windows with significant drug–baseline differences (cluster-based permutation, p<0.05, corrected), while light-gray bars indicate uncorrected significance for psilocybin (p < 0.05).

To further dissect network-specific temporal dynamics, we examined dReHo and dGFC trajectories across distinct cortical and subcortical networks for each psychedelic (Supplementary Figure S4-S8). During DMT exposure, dReHo decreased rapidly and persistently across nearly all functional networks except for LIM. In contrast, increases in dGFC were primarily confined to higher-order association networks (DMN and FPN), and the subcortical network. LSD induced widespread reductions in dReHo across all cortical and subcortical networks, with increased GFC emerging predominantly in the DMN, FPN, DAN, and VAN networks. Psilocybin also showed a similar trend, with widespread reductions in dReHo and broadly distributed increases in dGFC. Ketamine produced robust and sustained decreases in dReHo across cortical networks, with significant reductions persisting through much of the drug phase. In contrast, dGFC increases were more transient, with shorter periods of significance observed across cortical and subcortical networks. For nitrous oxide, dReHo reductions were observed across nearly all cortical networks except the LIM network, while dGFC increases were largely restricted to association networks such as the DMN and FPN. Together, these results demonstrate that psychedelics induce dynamically dissociable changes in ReHo and GFC, characterized by sustained and widespread reductions in ReHo alongside more selective and variable increases in GFC.

### Overlap of ReHo reductions across psychedelics

All five psychedelic compounds induced reductions in ReHo across widespread cortical and subcortical territories. However, the spatial extent and topography of these effects varied across compounds. To assess anatomical convergence in local desynchronization, we overlaid voxel-wise thresholded ReHo maps for each drug (Figure 5). Areas of overlap—primarily localized to transmodal regions such as the prefrontal cortex and temporoparietal junction (TPJ)—indicate common targets of psychedelic modulation. Notably, classical psychedelics (LSD, DMT, and psilocybin) and non-classical agents (ketamine and nitrous oxide) produced spatially consistent but distinct patterns of ReHo reductions.

**Figure 5:**
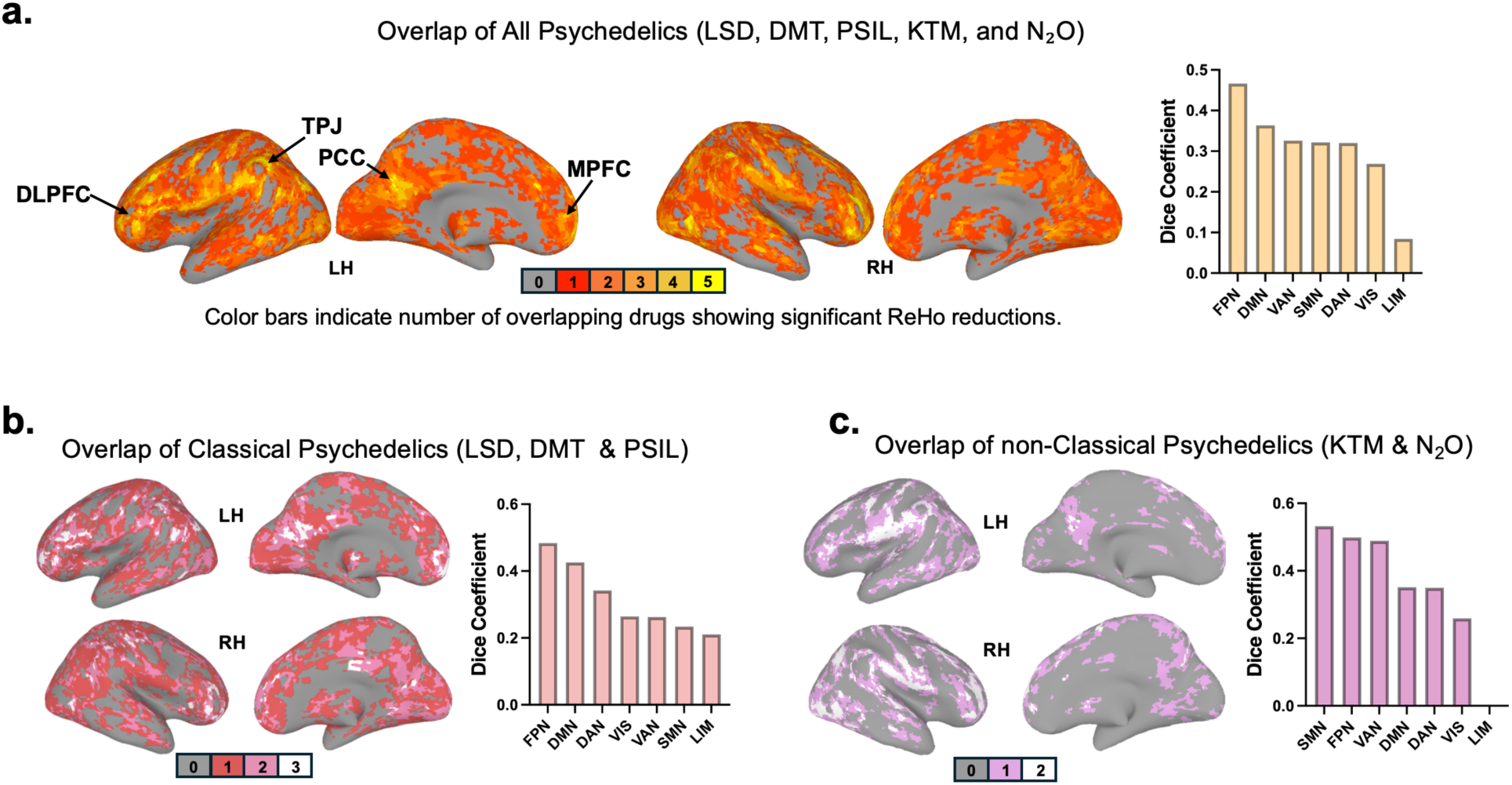
Overlap of regional homogeneity (ReHo) reductions across psychedelics. Voxel-wise thresholded ReHo reduction maps (p < 0.05, FWE-corrected) for each drug were overlaid to assess spatial convergence. **a.** overlap of all five psychedelics (dimethyltryptamine/DMT, lysergic acid diethylamide/LSD, psilocybin/PSIL, ketamine/KTM, and nitrous oxide/N_2_O). **b.** overlap between classical psychedelics (DMT, LSD, and psilocybin). **c.** overlap between non-classical psychedelics (ketamine and nitrous oxide). Color bars indicate the number of overlapping drugs showing significant ReHo reductions. Bar plots in each panel show the Dice coefficients summarizing the spatial overlap of ReHo reductions between psychedelic compounds within seven canonical cortical networks (DMN, FPN, DAN, VAN, SMN, VIS, LIM). Higher Dice values indicate greater overlap in the spatial distribution of ReHo reductions across drugs. Higher Dice values indicate greater overlap in the spatial distribution of ReHo reductions across drugs.

To quantify the degree of similarity among drug-induced ReHo change maps, we calculated global Dice coefficients, which quantify the degree of spatial overlap between thresholded statistical maps (ranging from 0 for no overlap to 1 for complete overlap) across the whole cortex. At the network level, overlap was most pronounced in large-scale association networks, particularly the FPN and DMN, whereas primary sensory networks showed weaker overlap. The LIM network consistently exhibited minimal overlap (Dice < 0.1 across all drug pairs), possibly reflecting both reduced signal sensitivity and genuine heterogeneity of drug effects within this system.

Together, these results indicate that psychedelic-induced ReHo reductions show partially overlapping yet compound-specific spatial profiles. The overlap is strongest in higher-order association networks, implying the convergence of psychedelic effects to a shared disruption of transmodal circuits across drugs of different pharmacological classes.

### Molecular and cellular predictors of ReHo reductions

Given the compound-specific spatial profiles of ReHo reductions observed across the five psychedelics, we next asked whether these divergent patterns could be explained by differences in underlying molecular and cellular architecture. To address this, we assembled a comprehensive set of molecular and cellular predictors. Neurotransmitter receptor predictors were taken from a high-resolution PET meta-atlas built from more than 1,200 healthy individuals across 19 receptors and transporters, providing reproducible, population-level estimates validated across tracers and cohorts ^31^. Cell-type predictors were derived from the Allen Human Brain Atlas using a transcriptomic modeling framework that generates spatially resolved maps of neuronal and non-neuronal cell classes ^32^ .

First, for each compound we performed multiple linear regression analyses relating ROI-wise ReHo changes (Cohen’s *d* across 450 regions) to standardized maps of neurotransmitter receptor and cell-type densities. To correct for spatial autocorrelation, we then implemented spin permutation testing with 10,000 surface-based rotations, generating empirical null distributions for adjusted R² values. All models yielded significant fits (FDR-corrected *p* < 0.05), suggesting that the spatial topographies of psychedelic-induced desynchronization are meaningfully constrained by neurobiological gradients at the molecular and cellular level.

Further, we then applied dominance analysis to rank each predictor based on its unique contribution to model variance. This approach decomposes the regression R² by estimating each predictor’s average marginal contribution across all possible submodels, enabling an interpretable hierarchy of molecular and cellular drivers. This revealed pharmacologically specific profiles:

ReHo reductions with classical psychedelics (DMT, LSD, and psilocybin) were most strongly associated with serotonergic features, especially 5-HT₂A, 5-HT₁A/₁B receptor densities. In contrast, the ReHo patterns with ketamine and nitrous oxide were most strongly associated with NMDA receptor densities (Figure 6). To directly compare dominance results between local (ReHo) and global (GFC) metrics, we additionally summarized the proportional contributions of receptor predictors in heatmaps (Supplementary Figure S9).

**Figure 6:**
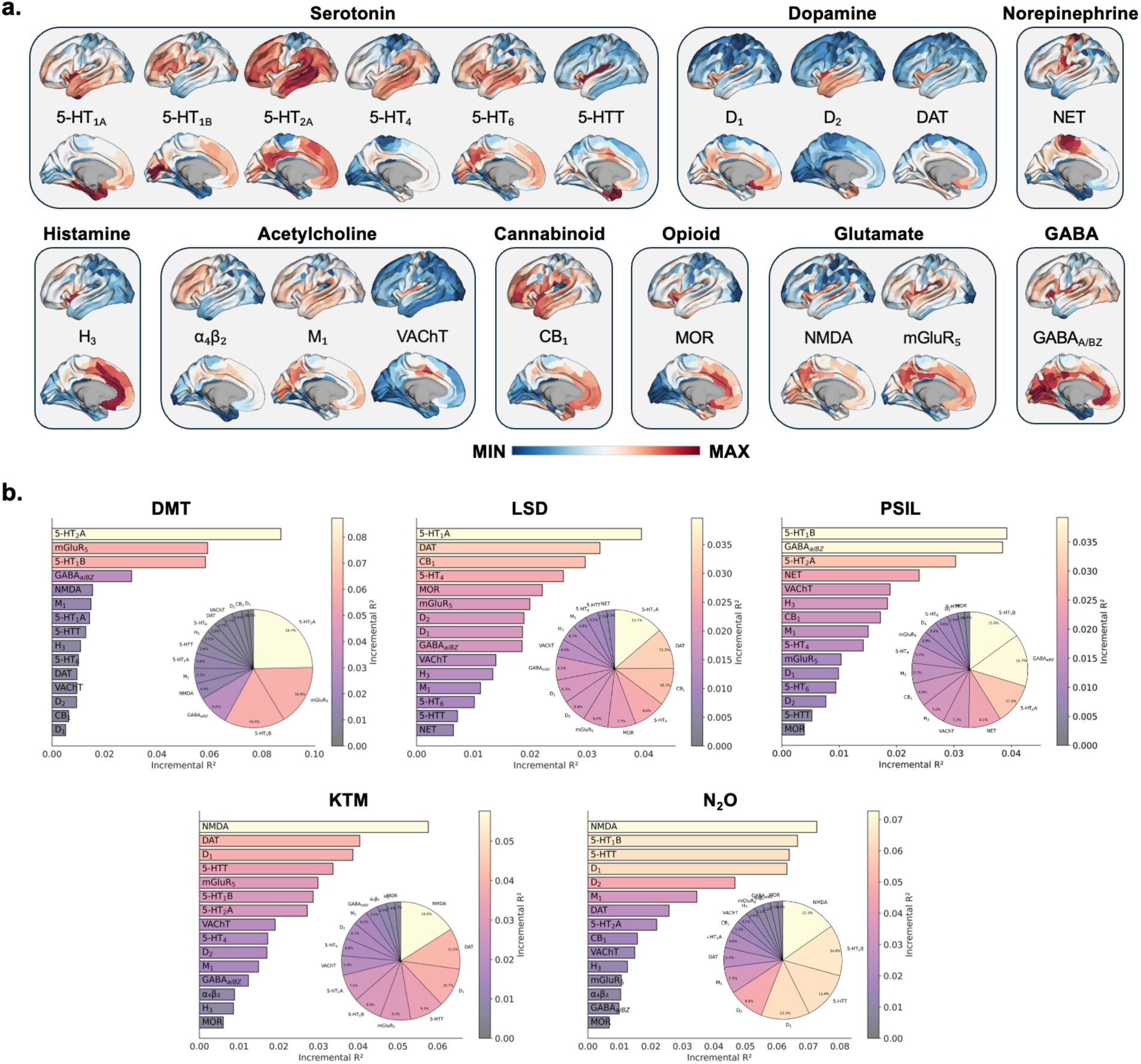
Dominance analysis of receptor density predictors of psychedelic-induced regional homogeneity (ReHo). **a.** Spatial distributions of receptor density maps derived from a high-resolution PET meta-atlas covering 19 neurotransmitter receptors and transporters across nine transmitter systems. Maps are z-scored across cortical regions (blue–red scale). **b.** Dominance analysis of receptor predictors. Bar plots depict the incremental R² values of each receptor density profile in explaining regional ReHo changes for dimethyltryptamine/DMT, lysergic acid diethylamide/LSD, psilocybin/PSIL, ketamine/KTM, and nitrous oxide/N_2_O. Pie charts summarize the proportional contributions of predictors for each compound. Abbreviations: 5-HT = serotonin; 5-HT₁A/B/2A/4/6 = serotonin receptor subtypes; 5-HTT = serotonin transporter; α₄β₂ = nicotinic acetylcholine receptor; M₁ = muscarinic acetylcholine receptor M1; VAChT = vesicular acetylcholine transporter; CB₁ = cannabinoid receptor type 1; D₁/D₂ = dopamine receptors; DAT = dopamine transporter; GABAₐ/BZ = GABA-A receptor/benzodiazepine site; H₃ = histamine H3 receptor; mGluR₅ = metabotropic glutamate receptor 5; MOR = μ-opioid receptor; NET = norepinephrine transporter; NMDA = N-methyl-D-aspartate receptor.

Across all five psychedelics, ReHo reductions were most strongly predicted by excitatory intratelencephalic (IT) projection neurons and MGE-derived inhibitory interneurons, including chandelier and somatostatin-expressing (Sst Chodl) cell classes. These two cell families consistently ranked among the top predictors across compounds, despite substantial differences in pharmacology. This convergence suggests that psychedelics may act through a shared cellular pathway involving perturbations to excitatory–inhibitory microcircuitry: increased engagement of IT neurons and reduced contribution from inhibitory MGE interneurons would shift local circuits toward greater excitation and diminished inhibitory control, producing the robust local desynchronization captured by ReHo. Notably, this pattern was evident across classical (DMT, LSD, psilocybin) and non-classical (ketamine, nitrous oxide) compounds, pointing to a common mesoscale mechanism underlying psychedelic-induced decreases in local synchrony. (Figure 7). To directly compare dominance results between local (ReHo) and global (GFC) metrics, we summarized the proportional contributions of cell type predictors in heatmaps (Supplementary Figure S10).

**Figure 7:**
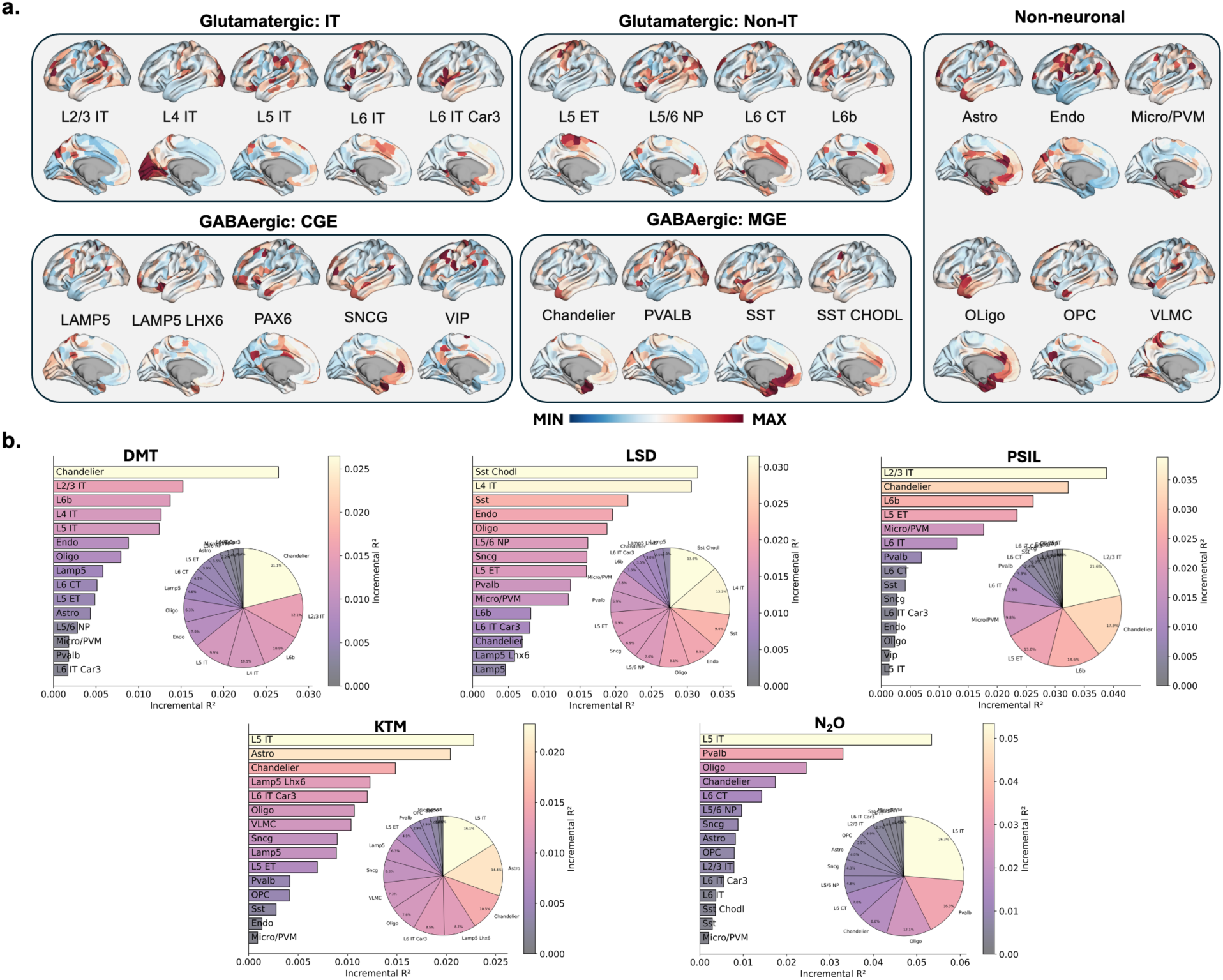
Dominance analysis of cell type density predictors of psychedelic-induced regional homogeneity (ReHo). **a.** Spatial distributions of modeled cell-type classes. Cortical cell-type density maps were derived from post-mortem transcriptomic data and grouped into major classes following the Allen Human Brain Atlas–based taxonomy, including glutamatergic intratelencephalic (IT) and non-IT projection neurons, GABAergic interneurons from the caudal ganglionic eminence (CGE) and medial ganglionic eminence (MGE) lineages, and major non-neuronal cell classes. **b.** Dominance analysis of cell-type predictors. Maps show z-scored relative density across the cortical surface (blue–red scale).Bar plots show the incremental variance explained (incremental R²) by each cell type density map in predicting region-wise reductions in regional homogeneity (ReHo) for dimethyltryptamine/DMT, lysergic acid diethylamide/LSD, psilocybin/PSIL, ketamine/KTM, and nitrous oxide/N_2_O. Pie charts summarize the relative percentage contributions of all predictors for each compound, derived from dominance analysis. Abbreviations: Lamp5 = Lamp5 interneurons; Lamp5 Lhx6 = Lamp5 Lhx6-positive interneurons; Pax6 = Pax6-expressing interneurons; Vip = VIP-positive interneurons; Sncg = Sncg-expressing interneurons; L5 ET = layer 5 extratelencephalic neurons; L5/6 NP = layer 5/6 near-projecting neurons; L6 CT = layer 6 corticothalamic neurons; L6b = layer 6b neurons; L2/3–L6 IT = layer-specific intratelencephalic neurons (Car3 = layer 6 IT Car3 subtype); Chandelier = chandelier cells; Pvalb = parvalbumin interneurons; Sst = somatostatin interneurons; Sst-Chodl = somatostatin Chodl-positive interneurons; Astro = astrocytes; Endo = endothelial cells; Micro/PVM = microglia/perivascular macrophages; Oligo = oligodendrocytes; OPC = oligodendrocyte precursor cells; VLMC = vascular and leptomeningeal cells.

### Linking receptor density, local desynchronization, and global connectivity

To bridge molecular targets with large-scale network integration, we conducted mediation analyses testing whether receptor density influences GFC indirectly via ReHo (Figure 8). We focused on the strongest receptor predictors identified in our dominance analyses for each compound. For DMT, 5-HT₂A receptor density significantly predicted stronger ReHo reductions, which in turn mediated increases in GFC (indirect effect: β = 0.083, p < 0.001). The direct path was not significant, indicating a fully mediated effect of serotonergic architecture on large-scale integration. For LSD, 5-HT₁A receptor density emerged as the dominant predictor, but neither the indirect pathway via ReHo nor the direct effect on GFC reached significance. For psilocybin, 5-HT₁B receptor density significantly predicted ReHo decreases, which in turn partially mediated increases in GFC (indirect β = −0.004, p = 0.016), consistent with a mixed pathway involving both local and global levels of organization.

**Figure 8:**
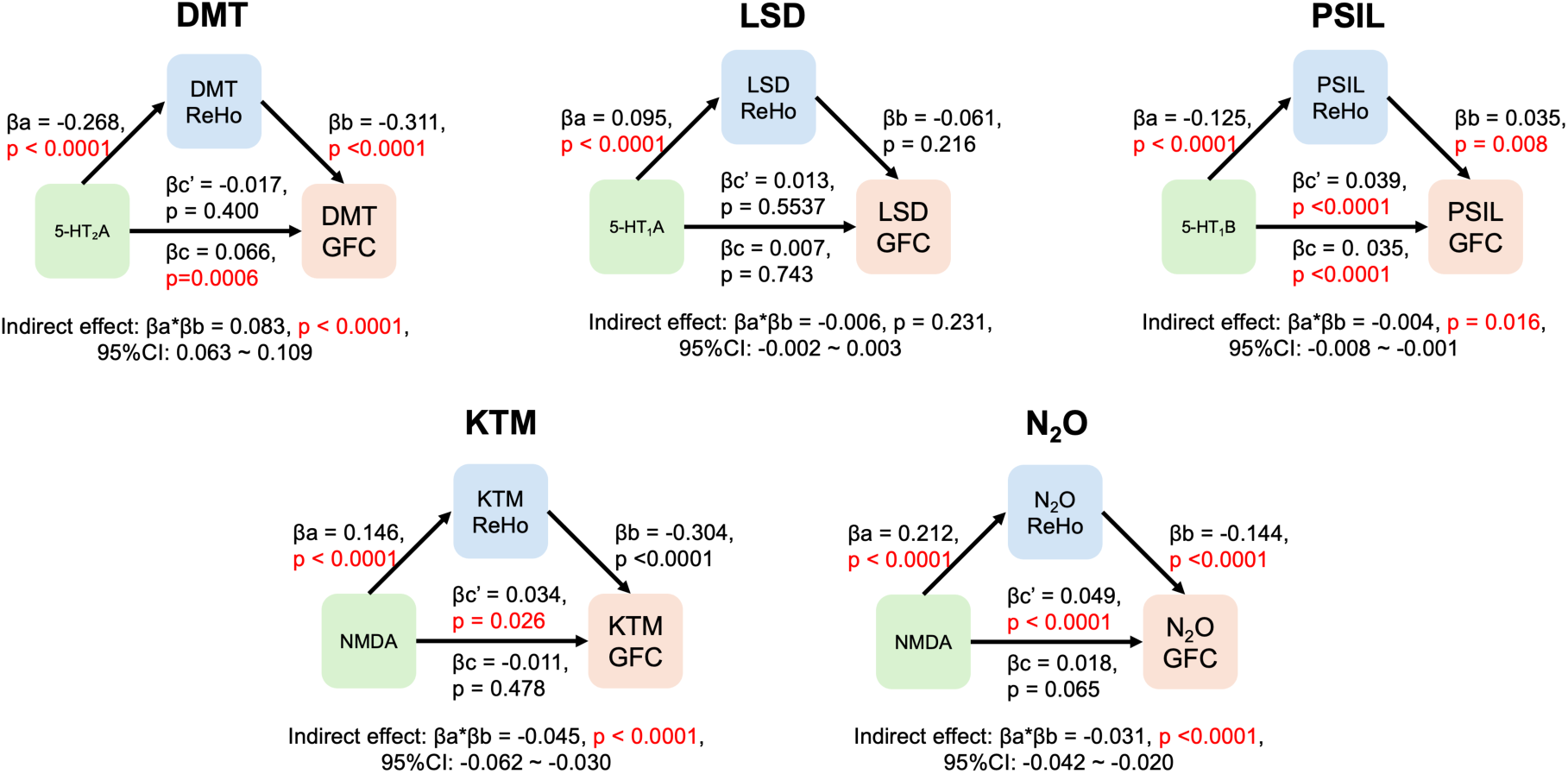
Mediation analyses linking receptor density, local synchrony (ReHo), and global connectivity (GFC) across five psychedelics. Mediation models tested whether receptor density (green box, X) predicted changes in global functional connectivity (orange box, Y) either directly (c′ path) or indirectly via changes in regional homogeneity (blue box, M; a × b path). For each compound (dimethyltryptamine/DMT, lysergic acid diethylamide/LSD, psilocybin/PSIL, ketamine/KTM, and nitrous oxide/N_2_O), standardized regression coefficients (β) and associated p-values are shown for each path, together with the estimated indirect effect (βa × βb) and 95% bootstrap confidence intervals (10,000 iterations).

In contrast, both ketamine and nitrous oxide showed significant mediation by NMDA receptor density. For ketamine, NMDA density predicted stronger local desynchronization, which in turn mediated changes in GFC (indirect β = −0.045, p < 1×10⁻⁸). For nitrous oxide, a similar NMDA-mediated pathway emerged (indirect β = −0.030, p < 1×10⁻⁹).

Together, these results delineate compound-specific, receptor–circuit–network relationships: for DMT, 5-HT₂A receptor density exerted its effects on global integration through ReHo-mediated desynchronization; for psilocybin, 5-HT₁B showed a weak but significant indirect effect via ReHo; in contrast, LSD showed no significant mediation, indicating that its serotonergic influences on GFC may operate independently of local synchrony. Non-classical agents converge on an NMDA-dependent mechanism where local desynchronization is a critical intermediary step.

## Discussion

In this study, we provide a multilevel characterization of how classical (DMT, LSD, and psilocybin) and non-classical (ketamine, nitrous oxide) psychedelics affect human brain function at the mesoscale. Despite distinct molecular targets, all five compounds consistently reduced regional cortical synchrony, particularly in higher-order association networks such as the default mode and frontoparietal systems. These local disruptions were most evident at fine spatial resolutions, demonstrating that psychedelic action engages microcircuit-level synchrony. Dynamic analyses revealed that sustained and widespread decreases in ReHo co-occurred with transient and spatially limited increases in global functional connectivity, indicating a dissociation between local desynchronization and large-scale integration.

A key contribution of this work is the focus on the mesoscale level of brain organization, which has remained underexplored in psychedelic research. Most prior studies emphasized either molecular mechanisms in animal models (e.g., receptor-level signaling pathways) or large-scale network reconfigurations in humans, particularly increases in global connectivity and network disintegration ^18,19,21,22,33^. What has been missing is a systematic investigation of how psychedelics alter local neural synchrony and how the latter affects macroscale functional connectivity in the human brain. Our approach addresses this gap by leveraging the measure of regional homogeneity (ReHo) to bridge molecular perturbations with network-level changes. By systematically comparing five pharmacologically distinct compounds, we show that diverse receptor systems and cellular substrates converge on a common mesoscale signature of disrupted local synchrony, while preserving compound-specific spatial profiles. Notably, the strongest reductions were observed in transmodal association networks such as the DMN and FPN, consistent with prior reports of network-level psychedelic reorganization ^21,22,30,34^, and highlighting sensitivity of these regions to local desynchronization.

Importantly, ReHo has already been widely applied as a local measure in psychiatric populations^35–40^, and the highly consistent reductions we observed across five distinct compounds underscore its robustness and establish it as a solid and generalizable measure of psychedelic action. Moreover, the ReHo effects proved highly stable: analyses with and without global signal regression (GSR) yielded similar patterns, with GSR only slightly increasing statistical power. Because ReHo reflects local dynamics, controlling for shared global variance (via GSR) may help isolate genuine local effects. Together, this suggests that ReHo captures locally coordinated dynamics that are relatively resilient to global signal confounds, unlike global functional connectivity, which is notably sensitive to GSR. Furthermore, our scale-dependent analyses revealed that these effects were largest at fine-grained neighborhoods (7–27 voxels), reinforcing the interpretation of ReHo as a sensitive mesoscale index of psychedelic action and underscoring the importance of spatial resolution in probing local brain dynamics. This scale-specific pattern aligns closely with emerging evidence from connectome-harmonic analyses, which consistently show a redistribution of brain activity from low-frequency, spatially smooth harmonics toward higher-frequency, spatially complex modes during psychedelic states ^41–43^. Higher-frequency harmonics reflect finely differentiated and spatially fragmented activity patterns. Our finding of strong ReHo reductions at the smallest voxel neighborhoods provides a mechanistic explanation for this shift: diminished local synchrony destabilizes the cortex’s ability to sustain smooth, low-frequency modes and instead promotes the expression of high-frequency, spatially heterogeneous patterns.

Extending static analysis to dynamic trajectories, we found a dissociation between sustained local desynchronization and more variable global integration, an effect not previously reported in psychedelic neuroimaging. Prior dynamic studies have characterized psychedelic effects primarily at global or network level, reporting, for example, increased dynamic global connectivity during DMT exposure ^29^, increased repertoire of functional connectivity states during LSD or psilocybin^44,45^, dynamic network reorganization through changes in control energy^46^, and altered co-activation patterns during nitrous oxide exposure ^33^. Our results demonstrate that local synchrony exhibits sustained dynamic disruption, highlighting a mesoscale dimension of psychedelic action that complements these macroscale findings.

Crucially, our dominance analysis results reveal bridges across spatial scales, by providing mechanistic evidence for how molecular and cellular substrates may shape brain-wide functional changes. Both sets of predictors (PET-derived neurotransmitter maps and transcriptomic cell-type reconstructions) are based on large, well-validated population datasets and therefore offer reliable anatomical priors for interpreting psychedelic-induced ReHo disruptions. Receptor-level findings showed clear pharmacological specificity. Classical psychedelics were most strongly associated with serotonergic receptor distributions, with DMT linked most strongly to 5-HT₂A density, psilocybin showing its top association with 5-HT₁B, and LSD primarily associated with 5-HT₁A. In contrast, the non-classical psychedelics ketamine and nitrous oxide were most strongly associated with NMDA receptor density. Although 5-HT₂A is widely viewed as the canonical mechanism of classical psychedelics^8,9^, our results refine this picture. A key insight comes from recent neurovascular-coupling work, which demonstrates that 5-HT₂A agonism can strongly alter the hemodynamic response function and even decouple neuronal activity from BOLD-based functional connectivity ^47^. This distinction helps resolve an otherwise puzzling feature in our data: ReHo, which indexes fine-scale local synchrony and is minimally affected by vascular confounds, showed weak correspondence with 5-HT₂A under LSD, whereas GFC, a BOLD-based global measure more sensitive to vascular and HRF modulation, showed strong 5-HT₂A dominance. Together, these observations suggest that LSD’s 5-HT₂A effects may propagate primarily through vascular or neurovascular pathways, enhancing GFC more than ReHo, whereas DMT and psilocybin exert more direct neuronally mediated serotonergic influences on local synchrony. Our mediation analyses provided a complementary perspective on how receptor distributions relate to global functional connectivity. For DMT and psilocybin, serotonergic receptor densities (5-HT₂A and 5-HT₁B) influenced global connectivity indirectly through their effects on local synchrony, indicating that ReHo partially mediates the propagation of molecular effects to systems-level organization. In contrast, LSD showed no significant mediation—its serotonergic contributions to GFC appear to operate independently of local synchrony. For ketamine and nitrous oxide, NMDA receptor density contributed indirectly via ReHo reductions. Overall, these results suggest that the link between molecular architecture, local desynchronization, and global connectivity varies systematically across psychedelic classes.

Beyond receptor-level contributions, our cell-type dominance analysis showed that ReHo reductions were most strongly associated with IT excitatory neurons and MGE-lineage inhibitory interneurons (PV, SST, chandelier), pointing to a disruption of local excitation–inhibition (E/I) balance as a core mesoscale mechanism. This interpretation aligns closely with electrophysiological findings: psychedelics reliably increase neural signal diversity (LZc) in MEG and EEG recordings^48^, a temporal signature of reduced local synchrony and destabilized microcircuit dynamics. In parallel, connectome-harmonic analyses^41^ show a shift from low-frequency, spatially smooth modes toward higher-frequency, spatially complex patterns during psychedelic states. The strong fine-scale ReHo reductions observed here provide a spatial correlate of this shift, suggesting that altered E/I microcircuitry promotes fragmentation of local dynamics, thereby enabling both increased temporal complexity and greater expression of high-frequency harmonic modes across the cortex.

In addition to advancing mechanistic understanding, our findings also have clinical relevance. Classic and atypical psychedelics—including LSD, DMT, psilocybin, ketamine, and nitrous oxide—have shown promise as rapid-acting interventions for depression and other psychiatric conditions^1,5,7,49^, yet neuroimaging measures that can guide their therapeutic use remain limited.

ReHo has already been widely employed in studies of psychiatric disorders such as depression and other mental illnesses, where changes in local synchrony are commonly observed in the patients^35–39^, as well as in neurodevelopmental conditions such as autism^50^. By demonstrating consistent reductions across multiple psychedelic compounds, our study helps align psychiatric neuroimaging research with emerging psychedelic therapy, suggesting a convergent neural mechanism across these domains. Moreover, because ReHo is a local measure, it enables spatially specific targeting of brain regions implicated in psychedelic effects. For example, we observed convergent reductions in the temporoparietal junction (TPJ), a densely connected multimodal associative hub linked to multiple functions including social cognition, attention, and perception ^51^. Importantly it has been proposed the TPJ plays a key role in the sense of self, and self-other distinctions ^52^, and has thus been linked to out-of-body experiences ^53,54^. These findings suggest the wide-ranging alterations in experience induced by these compounds may be underpinned by disruptions in TPJ function, suggesting a common neural substrate across compounds. Such localization opens the possibility of modulating these circuits directly, for instance via neuromodulation, to retain therapeutic efficacy while minimizing psychoactive effects. In this way, our results highlight how measures of local synchrony can not only bridge molecular and network levels but also inform the development of mechanism-based interventions.

Several limitations of this study should be acknowledged. First, despite testing multiple compounds in independent cohorts, sample sizes were modest and may limit generalizability. Replication in larger, prospectively studied cohorts will be important. Second, ReHo is an indirect hemodynamic measure of local synchrony, and future multimodal work—such as simultaneous electrophysiology or receptor-specific PET—will be critical to validate its neurobiological basis. Third, our analyses focused on acute drug effects, leaving open the question of whether local desynchronization is a transient state marker or contributes to longer-term plasticity that underlies therapeutic benefit. Fourth, the datasets included were acquired with different scanners and protocols, which, despite convergent findings, introduces potential variability that should be addressed through prospective, standardized designs. Fifth, we restricted our investigation to the neurobiological effects of psychedelics and did not correlate changes with psychedelic phenomenology. Linking the multiscale neural effects to subjective experiences will be critical for a comprehensive understanding. Sixth, although mediation analysis provided insights into the potential pathways linking receptor architecture, local synchrony, and global connectivity, such models cannot rule out latent or unmeasured variables. This limitation is inherent to causal statistical approaches, including structural path models. Finally, while we identified convergent reductions across five compounds, inter-individual variability in the magnitude and spatial distribution of ReHo changes remains an important direction for understanding predictors of subjective and clinical outcomes.

## Conclusions

Psychedelics consistently reduce local neural synchrony in the human brain, especially in higher-order association networks, while variably enhancing global connectivity. This common mesoscale signature links distinct receptor pathways to large-scale brain dynamics, suggesting that alterations in ReHo reflect a shared neural mechanism underlying psychedelic effects.

## Methods

### Subjects

#### Dataset 1: DMT

This study was approved by the National Research Ethics Committee London—Brent and the UK Health Research Authority and conducted in accordance with the Declaration of Helsinki (2000), International Council for Harmonization Good Clinical Practices guidelines, and the NHS Research Governance Framework. It was sponsored by Imperial College London and conducted under a Home Office license for Schedule 1 drug research. All participants gave written informed consent. More detailed description can be found in previous published work ^29^.

A total of 20 healthy participants (seven females; mean age ± standard deviation: 33.5 ± 7.9 years) completed all sessions. Exclusion criteria included: being under 18 years of age, contraindications for MRI, no prior psychedelic experience, prior adverse reaction to psychedelics, current or past significant psychiatric or physical illness (e.g., epilepsy, diabetes, heart disease), family history of psychosis, and excessive use of alcohol or other drugs of abuse.

The study employed a single-blind, placebo-controlled, within-subject, counterbalanced design. Each participant attended two sessions, spaced two weeks apart, at the Imperial College Clinical Imaging Facility. On one day, they received a placebo (10 mL sterile saline), and on the other, an intravenous dose of 20 mg DMT fumarate (dissolved in 10 mL sterile saline), administered over 30 seconds and followed by a 10 mL saline flush over 15 seconds. Drug administration occurred during the resting-state fMRI scan, at minute 8 of a continuous 28-minute session. Scanning continued for 20 minutes after injection. Participants rested in the scanner with eyes closed (a blindfold was used to ensure this), and simultaneous EEG was recorded.

Subjective drug effects were assessed using validated questionnaires administered after scanning, including the 11-Dimensional Altered States of Consciousness Questionnaire (11D-ASC) ^55^ and the Mystical Experience Questionnaire (MEQ-30). Additional scan sessions, where participants verbally rated drug intensity every minute in real time, were also conducted, but only the resting-state fMRI scans without in-scan ratings were analyzed in the present study. Intensity ratings from the cued sessions were used as covariates in dynamic fMRI and EEG analyses reported elsewhere.

fMRI data were acquired using a Siemens Magnetom Verio 3T MRI scanner (syngo MR B17) equipped with a 12-channel head coil compatible with electroencephalography. Functional images were collected using a T2*-weighted echo-planar imaging sequence with the following parameters: repetition time = 2000 ms, echo time = 30 ms, flip angle = 80°, voxel size = 3.0 × 3.0 × 3.0 mm³, 35 axial slices with no inter-slice gap, and total acquisition duration of 28 minutes. High-resolution T1-weighted structural scans were also acquired for registration purposes.

#### Dataset 2: LSD

This study was approved by the National Research Ethics Service London—West London Research Ethics Committee (14/LO/1190) and conducted in accordance with the Declaration of Helsinki. All participants provided written informed consent prior to participation. Further methodological details are available in the original reports ^22,56^.

Due to anxiety leading to an incomplete BOLD scan, one participant withdrew from the study, and four were excluded from group analysis owing to significant head motion. Consequently, 15 participants (including 5 females, with a mean age and standard deviation of 38.4±8.6 years) were retained in the dataset.

Exclusion criteria encompassed a history of drug abuse and psychosis, among other health-related conditions detailed in the original reports. The study design included two sessions where participants received either a placebo or LSD in a counterbalanced order. A medical professional inserted and secured a cannula in the antecubital fossa vein of each volunteer. Each participant was administered 75 μg of LSD intravenously, using a 10 ml solution infused over two minutes, followed by a saline infusion. MRI scans commenced around 70 minutes post-dosing to observe peak intensity alterations occurring between 60 and 90 minutes following administration.

The imaging process was conducted using a 3T GE HDx system. For capturing functional images of the entire brain, a gradient-echo echo-planar imaging pulse sequence was employed. The specific parameters set for this sequence were as follows: 35 slices, a repetition time and echo time of 2000/35 milliseconds, a slice thickness of 3.4 millimeters, a field of view extending 220 millimeters, an image matrix sized at 64 × 64, a flip angle established at 90 degrees, and the total scan duration was 7 minutes. In addition, high-resolution anatomical images were acquired.

#### Dataset 3: Psilocybin

This dataset was collected at Washington University in St. Louis as part of a randomized cross-over precision functional brain mapping study (ClinicalTrials.gov identifier NCT03866174). The protocol was approved by the Washington University Institutional Review Board, and all participants provided written informed consent. Psilocybin was manufactured under good clinical practice by the Usona Institute, and all dosing sessions were facilitated by two trained clinical research staff to ensure participant safety. Further methodological details are available in the original report^57^.

Seven healthy adults (aged 18–45 years) were enrolled between 2021 and 2023. All participants had prior lifetime psychedelic exposure but none within the 6 months preceding the drug exposure. Exclusion criteria included psychiatric illness (depression, psychosis, or addiction), drug abuse, or contraindications for MRI. One participant could not tolerate psilocybin scanning and was excluded from group analysis, leaving six participants for inclusion.

The study employed a cross-over design with both psilocybin and an active placebo control (40 mg methylphenidate). Each participant completed multiple non-drug baseline sessions, followed by randomized drug sessions (25 mg oral psilocybin or 40 mg methylphenidate), with MRI commencing 60–180 minutes post-dosing to coincide with peak drug effects. Replication sessions were conducted 6–12 months later, including additional baseline and psilocybin imaging.

Imaging was performed on a Siemens Prisma 3T MRI scanner. Resting-state fMRI scans were acquired with a multiband, multi-echo echo-planar imaging sequence (voxel size = 2.0 mm isotropic; TR = 1761 ms; TEs = 14.2, 38.9, 63.7, 88.4, 113.1 ms; flip angle = 68°; multiband factor = 6; in-plane acceleration = 2; 72 axial slices; 510 frames; duration = 15 min 49 s). Each session included two resting-state scans, and in some cases additional task-based fMRI was acquired. High-resolution T1- and T2-weighted anatomical scans were also collected at 0.9 mm isotropic resolution.

#### Dataset 4: Ketamine

The study received approval from the Institutional Review Board of Huashan Hospital, affiliated with Fudan University, and written informed consent was obtained from every participant. The research enrolled 12 right-handed individuals (including five females), with mean age and standard deviation of 41.4±8.6 years. These volunteers were classified as American Society of Anesthesiologists physical status I or II and had no recorded history of neurological dysfunction, significant organ impairment, or consumption of neuropsychiatric medication.

Administration of ketamine occurred via an intravenous line inserted into the left forearm vein. fMRI was continuously performed during the experiment, which lasted between 44 and 62 minutes, averaging 54.6 ± 5.9 minutes. Initially, a 10-minute baseline conscious state was established (excluding two subjects who had baseline periods of 6 and 11 minutes, respectively). Subsequently, a ketamine dose of 0.05 mg/kg per minute was administered for 10 minutes (totaling 0.5 mg/kg), followed by a 0.1 mg/kg per minute dosage for another 10 minutes (accumulating to 1.0 mg/kg in total), except for two participants, who received only the latter dosage. After discontinuation of the ketamine infusion, participants naturally regained connected consciousness. The entire procedure was supervised by two qualified anesthesiologists, with emergency resuscitation equipment readily accessible. Our analysis focused only on the subanesthetic ketamine levels.

A Siemens 3T MAGNETOM scanner, with a conventional eight-channel head coil, was utilized for the study. The whole-brain functional images were acquired using a gradient-echo echo-planar imaging pulse sequence. The parameters set for this sequence included: 33 slices, repetition time and echo time at 2000/30 milliseconds, a slice thickness of 5 millimeters, a field of view measuring 210 millimeters, an image matrix of 64 × 64, a flip angle of 90 degrees, and the scan duration was set at 12 minutes. Additionally, high-resolution anatomical images were obtained.

#### Dataset 5: Nitrous oxide

The investigation, approved by the University of Michigan Medical School’s IRB (HUM00096321), was part of a clinical trial (NCT03435055). Of the participants, written informed consent was obtained from each individual, two were excluded due to significant head motion (affecting 50% of their fMRI volumes), leaving a total of 16 subjects (8 females, mean age ± standard deviation: 24.6±3.7 years) who completed two fMRI resting-state scans before and during the administration of subanesthetic concentrations (35%) of nitrous oxide. All participants were categorized as American Society of Anesthesiologists physical status I, signifying good health. Exclusion criteria included a history of drug abuse and psychosis, among other health-related conditions (detailed in the published registry: https://www.clinicaltrials.gov/ct2/show/NCT03435055).

This study adopted a within-subject design for both fMRI and altered states assessment. Each subject, after completing an altered-states questionnaire, underwent fMRI during both placebo and 35% nitrous oxide inhalation. Nitrous oxide administration commenced pre-scan to ensure equilibrium. The focus of the current study was solely on resting-state fMRI data analysis.

MRI-compatible anesthesia machines ensured safety, with anesthetic administration overseen by expert anesthesiologists. fMRI data collection for the study was carried out at Michigan Medicine, University of Michigan, employing a Philips Achieva 3T scanner. Functional brain images were captured using a T2*-weighted echo-planar imaging sequence with specific parameters: 48 slices, repetition/echo times of 2000/30ms, 3 mm slice thickness, and a 200 × 200mm field of view. The flip angle was set at 90 degrees and each scan lasted 6 minutes. Additionally, high-resolution anatomical images were acquired to aid in co-registering with resting-state fMRI data.

#### fMRI data preprocessing

For the preprocessing of fMRI dataset 1, 3, 4 and 5 in this study, the AFNI software (available at http://afni.nimh.nih.gov/) was utilized. The procedure encompassed several steps: First, the initial two frames of each scan were removed to ensure signal stability. This was followed by slice timing correction to adjust for temporal differences in the acquisition of slices. Next, rigid head motion correction and realignment were performed. The frame-wise displacement (FD) for head motion was quantified as the Euclidean Norm of the six-dimensional motion derivatives. Any frame with a derivative value exceeding an FD of 0.4mm, along with its preceding frame, was excluded. The next step involved coregistration with T1 anatomical images for accurate alignment and integration. This was followed by spatial normalization into Talairach space^58^ and resampling to 3 mm isotropic voxels to standardize the spatial coordinates of the images. Time-censored data then underwent band-pass filtering in the range of 0.01–0.1Hz using AFNI’s function 3dTproject. Concurrently, linear regression was employed to remove undesired components from the data, including linear and nonlinear drift, time series of head motion and its temporal derivative, and mean time series from white matter and cerebrospinal fluid. Subsequently, spatial smoothing was applied using a 6mm full-width at half-maximum isotropic Gaussian kernel. Finally, normalization of each voxel’s time series to zero mean and unit variance was carried out, ensuring consistency and comparability across the dataset.

Dataset 2, referenced at doi:10.18112/openneuro.ds003059.v1.0.0, was processed through several steps. Initially, the first three volumes of each scan were removed for stability. De-spiking was then conducted to correct signal artifacts, followed by slice time correction to align image acquisition timings. Motion correction was applied to counteract participant movement, and brain extraction isolated brain tissue from other elements in the images. The images were aligned to anatomical scans via rigid body registration and further aligned to a 2mm MNI brain template through non-linear registration. The dataset was then scrubbed using a frame-wise displacement threshold of 0.4, with a maximum of 7.1% of volumes scrubbed per scan, replacing them with the mean of surrounding volumes. Additional steps included spatial smoothing with a 6mm kernel, band-pass filtering from 0.01 to 0.08 Hz, linear and quadratic de-trending to remove signal drifts, and regressing out undesired components related to motion and anatomy. These procedures were integral in ensuring the quality and reliability of the dataset for subsequent analyses.

#### Regional Homogeneity (ReHo) analysis

In our study, ReHo was computed to quantify the local synchronization of neural activity by examining the time-series similarity within clusters of neighboring voxels. The ReHo metric was derived using Kendall’s coefficient of concordance (Kendall’s W), which measures the degree of agreement among the ranks of time series data. The ReHo values for each voxel were calculated using the following formula:

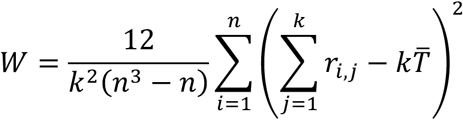

Here, *r_i,j_* signifies the rank of the ith time point in the jth voxel, 𝑘 represents the number of voxels in the cluster, 𝑛 is the number of time points, and 𝑇 is the mean rank across all voxels and time points. The value of Kendall’s W ranges from 0, indicating no agreement, to 1, indicating complete agreement, thus reflecting the degree of local functional connectivity. After the ReHo computation, we applied Fisher’s z-transformation to the ReHo values. This transformation standardizes the variance of the ReHo values, facilitating more accurate correlation studies and comparisons across different subjects and conditions. We also implemented a variation in the cluster size for the calculation of Regional Homogeneity. This variation was crucial to understand how different spatial extents of clusters influence ReHo measurements. Specifically, we chose cluster sizes corresponding to different Brain Radii settings (detailed in AFNI’s 3dReHo documentation). These settings correspond to clusters comprising varying numbers of voxels, namely 7, 19, 27, 25, 343, 729, 1331, 2197, 3375.

#### Overlap analysis of voxel-wise ReHo reductions across psychedelics

To assess spatial convergence of local desynchronization effects across psychedelic conditions, we conducted a voxel-wise overlap analysis based on statistical ReHo maps. For each drug (LSD, DMT, psilocybin, ketamine, and nitrous oxide), we first identified voxels showing significant ReHo reductions relative to baseline at p < 0.05, FWE-corrected. The resulting thresholded statistical maps were binarized such that significantly reduced voxels were assigned a value of 1, and non-significant voxels were set to 0. These binary maps were then summed across drugs to generate voxel-wise overlap maps, where each voxel’s value reflects the number of drugs showing significant ReHo reductions at that location. Separate maps were generated for all five drugs combined, classical psychedelics only (DMT, LSD, and psilocybin), and non-classical psychedelics only (ketamine and nitrous oxide). Overlap maps were visualized on inflated cortical surfaces and the color scale represents the number of overlapping drugs per voxel.

#### Global functional connectivity

To assess global functional connectivity (GFC), we first parcellated the brain into 400 cortical regions of interest (ROIs) using the Schaefer atlas. For each participant, we computed a 400 × 400 Pearson correlation matrix across all ROI time series. Fisher’s z-transformation was applied to normalize the correlation coefficients. The GFC value for each ROI was defined as the average of its Fisher z-transformed correlations with all other 399 ROIs in the brain, yielding a single GFC value per ROI. To derive network-level GFC, we then extracted the ROIs belonging to each of the seven Yeo functional networks ^59^ and averaged their corresponding ROI-level GFC values. This approach enabled the quantification of global connectivity at both regional and network scales.

#### Dynamic regional homogeneity and functional connectivity

To assess time-varying changes in local and global brain function, we computed dynamic ReHo and dynamic global functional connectivity using a sliding window approach. For each subject, a fixed-length sliding window of 60 repetition times with a step size of 1 repetition time was applied across the entire resting-state fMRI time series.

Within each window, ReHo was computed using the same method described above. The resulting voxel-wise ReHo maps were parcellated into 450 ROIs covering cortical and subcortical areas, and mean ReHo values were extracted for each ROI per window to obtain a dynamic time series.

For dynamic GFC, we computed a 450 × 450 ROI-wise Pearson correlation matrix within each window and applied Fisher’s z-transformation. For each ROI, GFC was defined as the average of its z-transformed correlations with all other ROIs. These ROI-level GFC values were then averaged across all ROIs to derive a whole-brain GFC time series. Network-level dynamic GFC was obtained by averaging ROI-level GFC values within each of the seven canonical Yeo networks.

To identify time periods showing significant condition-level differences, pairwise comparisons between DMT and placebo were conducted at each time point for both dynamic ReHo and dynamic GFC. This analysis was carried out at the whole-brain level and within each of the seven Yeo networks. Significant clusters of consecutive time points were identified using cluster-based permutation testing with 5,000 iterations, based on permutations of within-subject condition labels.

#### Neurobiological predictors and statistical modeling

Neurotransmitter receptor predictors were derived from a PET meta-atlas^31^, which integrates data from multiple independent cohorts to generate group-averaged tracer maps for 19 neurotransmitter receptors, transporters, and binding sites across nine major neurotransmitter systems (dopamine, norepinephrine, serotonin, acetylcholine, glutamate, GABA, histamine, cannabinoid, and opioid). For receptors with multiple tracers, the map based on the largest sample size was selected. All maps were resampled to template space and z-scored across voxels.

Cell-type predictors were obtained from the Allen Human Brain Atlas^32^ and included excitatory projection neurons (intratelencephalic, pyramidal tract, corticothalamic), inhibitory interneurons (parvalbumin-positive and chandelier cells), and glial classes (astrocytes, oligodendrocytes, microglia). Gene expression data were parcellated according to the same atlas used for fMRI analyses and z-scored for comparability.

For each psychedelic drug, ROI-wise ReHo changes (Cohen’s d across 450 ROIs) served as the dependent variable in a multiple linear regression framework, with receptor and cell-type densities as standardized predictors. To account for spatial autocorrelation, model significance was evaluated using a spin permutation test: functional change maps were randomly rotated on the cortical surface (10,000 iterations), and the regression model was recomputed at each iteration to generate a null distribution of adjusted R² values. Empirical p-values were derived by comparing observed model fits against this null distribution.

To further evaluate the relative importance of predictors, dominance analysis was applied. This variance-partitioning approach assesses predictor contributions across all possible submodels. Analyses were conducted separately for ReHo and GFC outcomes, with predictor variables z-scored prior to analysis. Normalized dominance values were reported, representing the proportion of variance explained by each predictor relative to the full-model R².

#### Mediation analysis

Mediation analysis was performed to test whether changes in local brain synchrony (ReHo) mediated the relationship between regional transmitter receptor density (X) and alterations in global functional connectivity (GFC; Y) for each drug condition.

For each compound, receptor/transporter density maps derived from PET atlases were entered as predictors (X), ReHo change values (drug – baseline; M) were treated as the mediator, and GFC change (drug – baseline; Y) was the outcome. The mediation model was specified as follows: Path a: effect of receptor density on ReHo (X → M). Path b: effect of ReHo on GFC, controlling for receptor density (M → Y). Path c′ (direct effect): direct effect of receptor density on GFC controlling for ReHo (X → Y). Path c (total effect): total effect of receptor density on GFC (X → Y, without M in the model). Indirect effect (ab): product of path a and path b, representing the mediated effect through ReHo.

All coefficients were estimated using the delta method for standard errors, and significance of indirect effects was assessed using percentile bootstrap confidence intervals (10,000 resamples). A significant indirect effect (ab) indicates the presence of mediation. We further classified mediation patterns into full mediation (ab significant, c′ not significant), partial mediation (ab and c′ both significant, same direction), and competitive mediation (ab and c′ significant but in opposite directions).

#### Statistical Analysis

Comparisons were made against placebo (intravenous saline) for DMT and LSD, and against the normal awake baseline for psilocybin, ketamine, and nitrous oxide.

For time average (static) ReHo, voxel-wise paired t-tests were conducted for DMT, LSD, ketamine, and nitrous oxide, with significance defined at p < 0.05, family-wise-error corrected. Network-level and whole-brain mean ReHo values were compared using paired t-tests with False Discovery Rate (FDR) correction (pFDR < 0.05). For psilocybin, static ReHo changes were assessed using linear mixed-effects models to account for repeated sessions.

For GFC, paired t-tests were applied at whole-brain and network levels, with results corrected using FDR (pFDR < 0.05).

For dynamic analyses (sliding-window ReHo and GFC), statistical significance was assessed with permutation-based cluster correction (5,000 iterations, p < 0.05). For psilocybin, dynamic ReHo and GFC trajectories were compared against baseline using paired t-tests.

For overlap/probability analyses, thresholded voxel-wise ReHo maps (p < 0.05, family-wise-error corrected) were overlaid across drugs, and spatial similarity was quantified using Dice coefficients at whole-cortex and network levels.

For dominance analysis, multiple linear regression models were fitted with receptor and cell-type densities as predictors of ReHo or GFC changes. Predictor importance was quantified using normalized dominance values, which divide the explained variance (R²) among predictors across all possible regression submodels, reflecting each predictor’s proportional contribution to the overall model.

For mediation analysis, we tested whether receptor density influenced GFC indirectly via ReHo. Indirect (mediated) effects were estimated using non-parametric bootstrap resampling (5,000 iterations). Mediation was considered significant if the 95% confidence interval of the indirect effect did not include zero.

## Data availability

Data used in the analyses will be publicly available from Zenodo repository. The LSD dataset is published at Openneuro (doi: 10.18112/openneuro.ds003059.v1.0.0).

## Code availability

Publicly available software and toolboxes used for analysis and visualization include AFNI (https://afni.nimh.nih.gov), JASP (v0.16.3; https://jasp-stats.org), MATLAB R2022a (https://www.mathworks.com), Prism (version 9.5.0; https://www.graphpad.com), and PyCharm (v2023.2.1; https://www.jetbrains.com/pycharm).

## Acknowledgements

This work was funded by National Institutes of Health (Bethesda, Maryland, USA) grants R01-GM103894 (to A.G.H. and Z.H.), R01-GM111293 (to G.A.M.) and T32-GM103730 (to G.A.M., PI, and R.D., Z.H., Fellows).

## Authorship contribution statement

Conceptualization: R.D., Z.H., G.A.M. Methodology: R.D., Z.H., G.A.M. Investigation: R.D., Z.H., R.C., G.A.M. DMT Data collection: C.T., R.L.C. Data analysis and visualization: R.D., Supervision: G.A.M, A.G.H., Writing—original draft: R.D., Writing—review & editing: All authors.

## Supplementary Material

**Figure S1:**
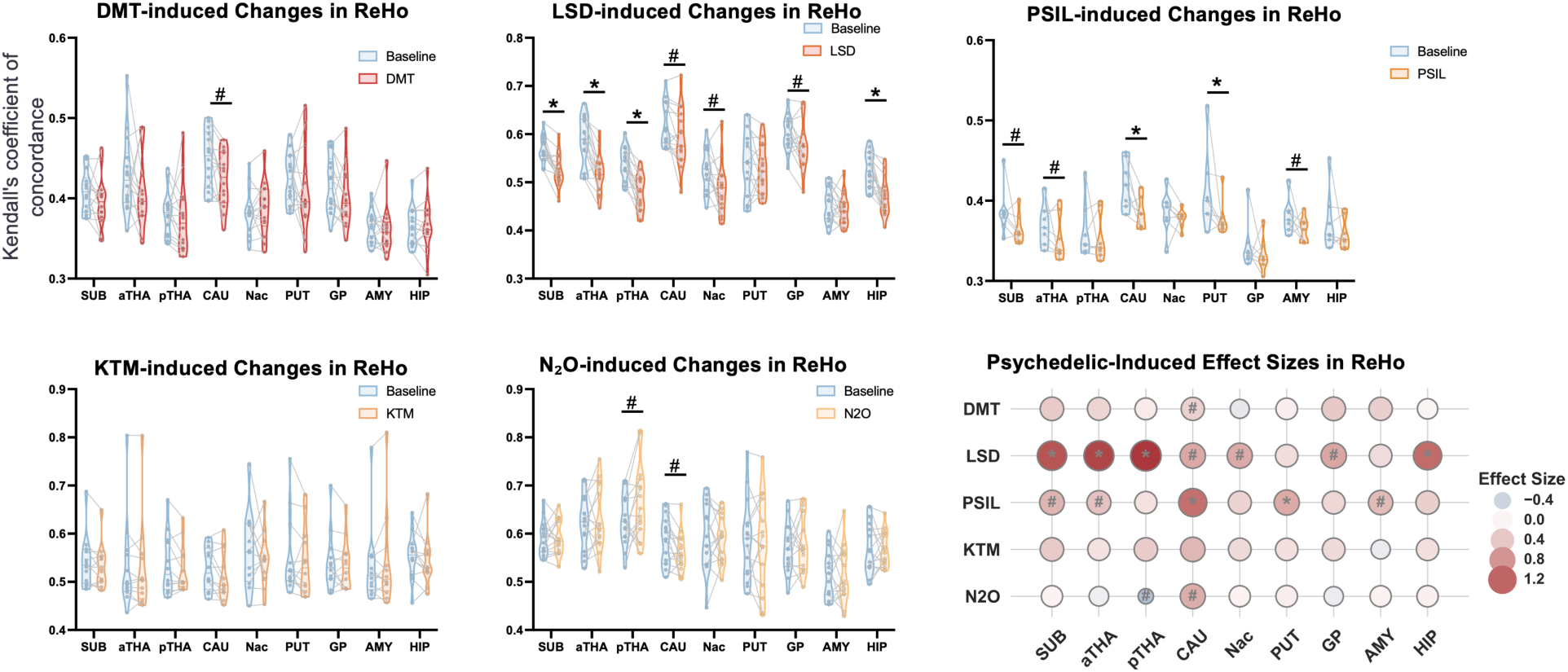
Subcortical regional homogeneity (ReHo) changes across psychedelic states. Violin plots (left) show ReHo values for the subcortical average (SUB) and individual regions (aTHA: anterior thalamus; pTHA: posterior thalamus; CAU: caudate; Nac: nucleus accumbens; PUT: putamen; GP: globus pallidus; AMY: amygdala; HIP: hippocampus) in each drug condition compared to baseline. Bubble plots (right) illustrate effect sizes (Cohen’s d) for baseline versus drug. Asterisks denote statistically significant reductions after FDR correction; number signs (#) indicate significance before correction. DMT: dimethyltryptamine, LSD: lysergic acid diethylamide, PSIL: psilocybin, KTM: ketamine, N_2_O: nitrous oxide.

**Figure S2:**
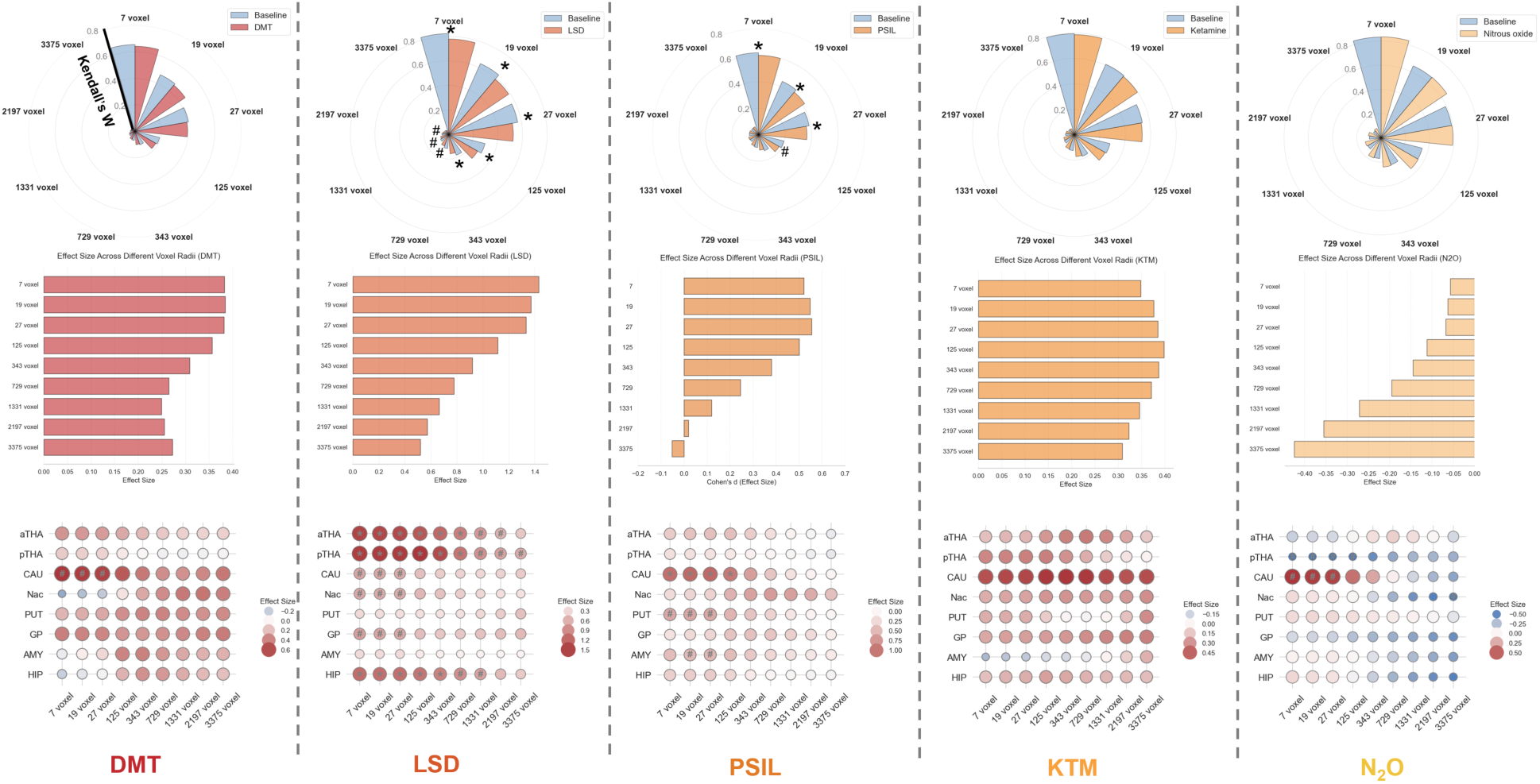
Spatial-scale dependence of psychedelic-induced ReHo changes in subcortical regions. For each drug (dimethyltryptamine/DMT, lysergic acid diethylamide/LSD, psilocybin/PSIL, ketamine/KTM, and nitrous oxide/N_2_O), regional homogeneity (ReHo) was computed using spherical neighborhoods from 7 voxels (radius = 1 voxel) to 3375 voxels (radius = 7 voxels). Polar plots (top) show subcortical average ReHo values for drug versus baseline across radii, with asterisks denoting significant reductions after FDR correction. Bar plots (middle) present effect sizes (Cohen’s d) of subcortical average ReHo changes across spatial scales. Bubble plots (bottom) depict effect sizes in individual subcortical regions (aTHA: anterior thalamus; pTHA: posterior thalamus; CAU: caudate; Nac: nucleus accumbens; PUT: putamen; GP: globus pallidus; AMY: amygdala; HIP: hippocampus) across different radii.

**Figure S3:**
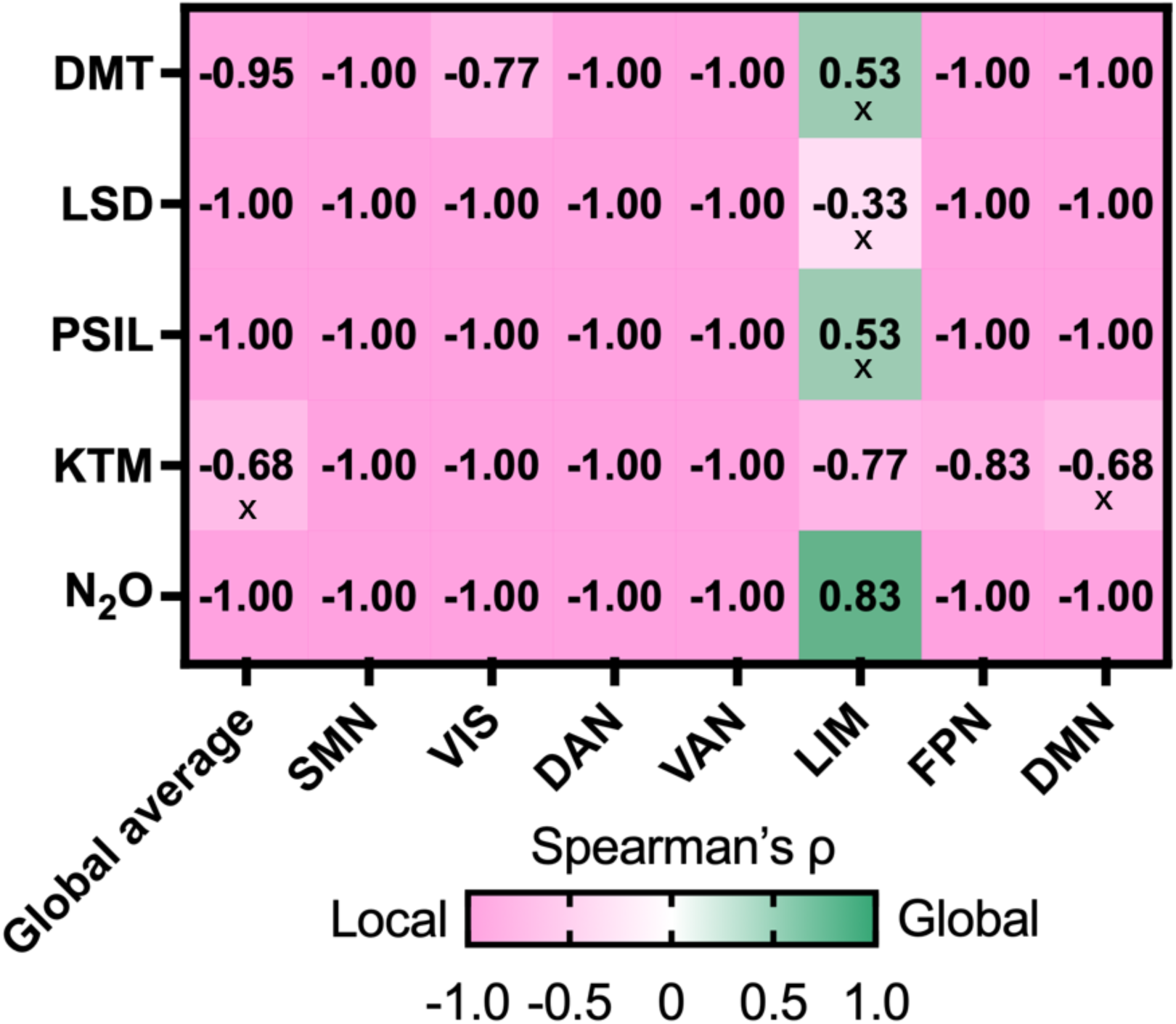
Spearman correlations between spatial scale and ReHo effect sizes. For each drug (rows) and cortical network (columns; global average plus seven canonical networks), we computed Spearman’s ρ between neighborhood radius (voxel size / voxel count across scales) and the corresponding ReHo effect sizes (Cohen’s d). Negative values indicate stronger ReHo reductions at finer spatial scales, whereas positive values indicate relatively larger effects at coarser scales. Color encodes the direction and magnitude of the scale–effect relationship (pink = more local/fine-scale dominance; green = more global/coarse-scale dominance). Cross marks indicate correlations that did not survive FDR correction.

**Figure S4:**
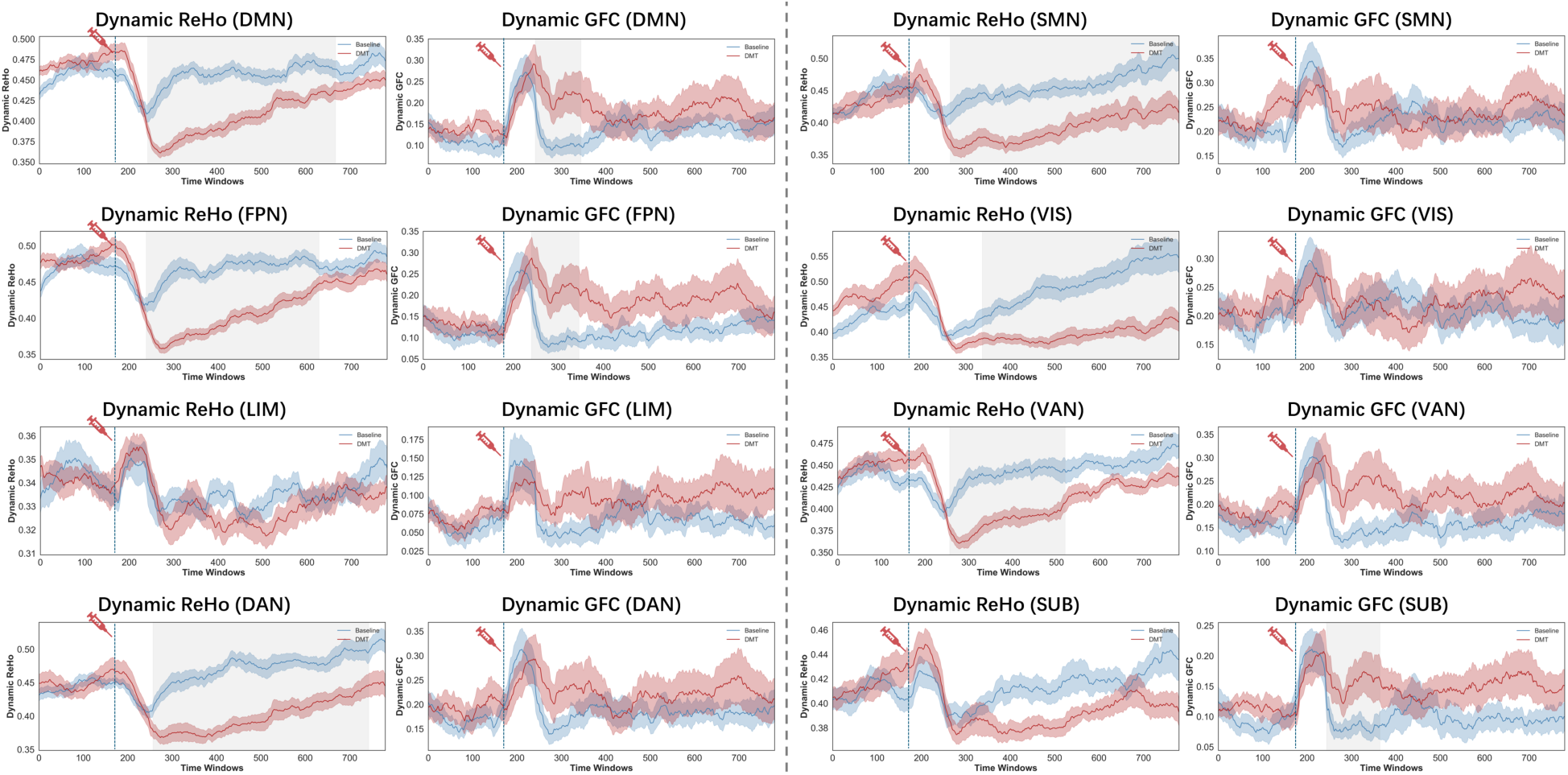
Dynamic regional homogeneity (ReHo) and global functional connectivity (GFC) in the DMT condition. Dynamic ReHo (left) and dynamic GFC (right) were computed using a sliding-window approach (window = 60 TRs, step = 1 TR) across the resting-state fMRI time series. Solid lines represent group mean values, with shaded bands indicating the standard error. Gray bars beneath each time course denote consecutive time windows where drug–baseline differences reached significance, as determined by cluster-based permutation testing (5,000 iterations).

**Figure S5:**
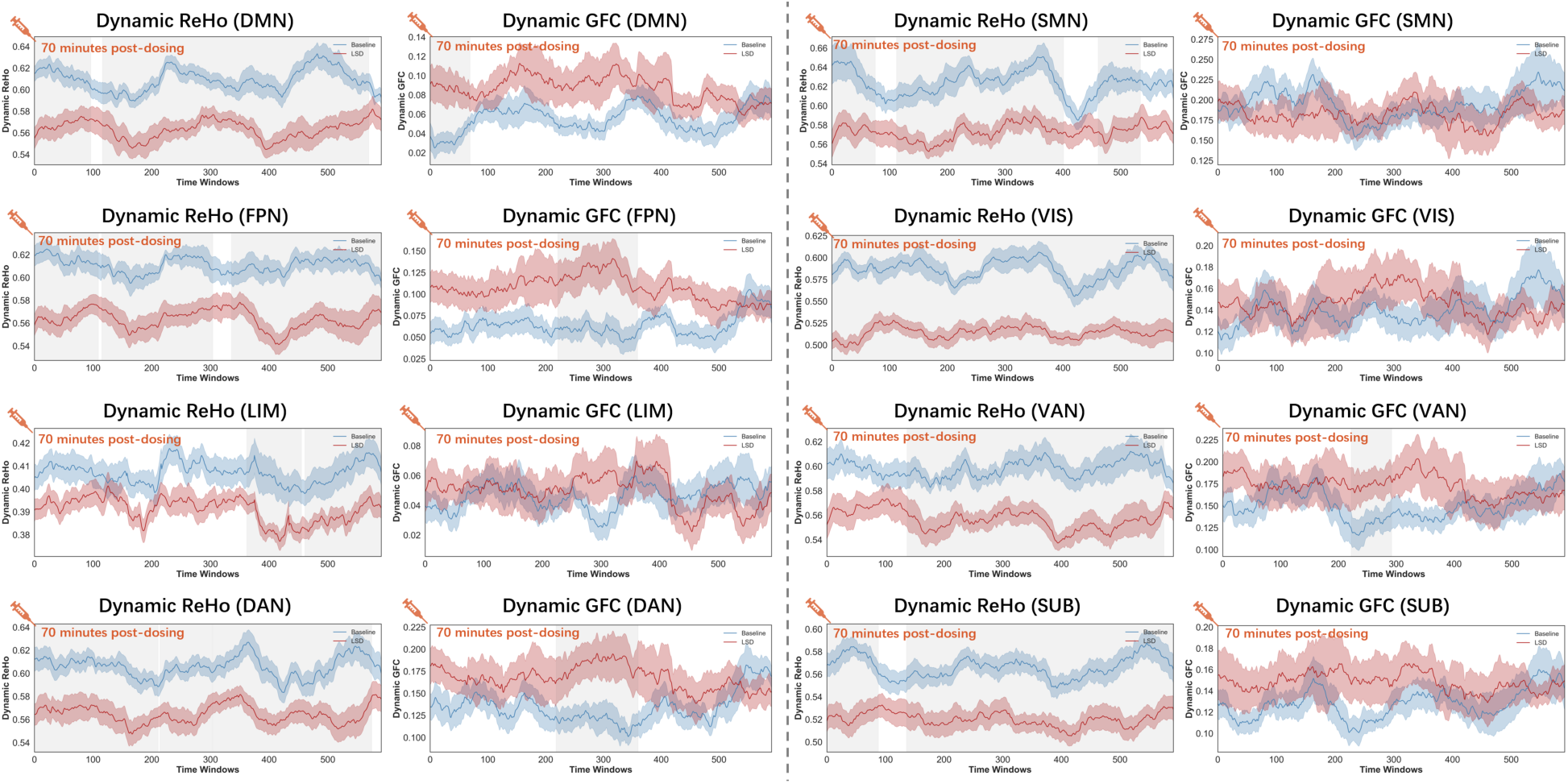
Dynamic regional homogeneity (ReHo) and global functional connectivity (GFC) in the LSD condition. Dynamic ReHo (left) and dynamic GFC (right) were computed using a sliding-window approach (window = 60 TRs, step = 1 TR) across the resting-state fMRI time series. Solid lines represent group mean values, with shaded bands indicating the standard error. Gray bars beneath each time course denote consecutive time windows where drug–baseline differences reached significance, as determined by cluster-based permutation testing (5,000 iterations).

**Figure S6:**
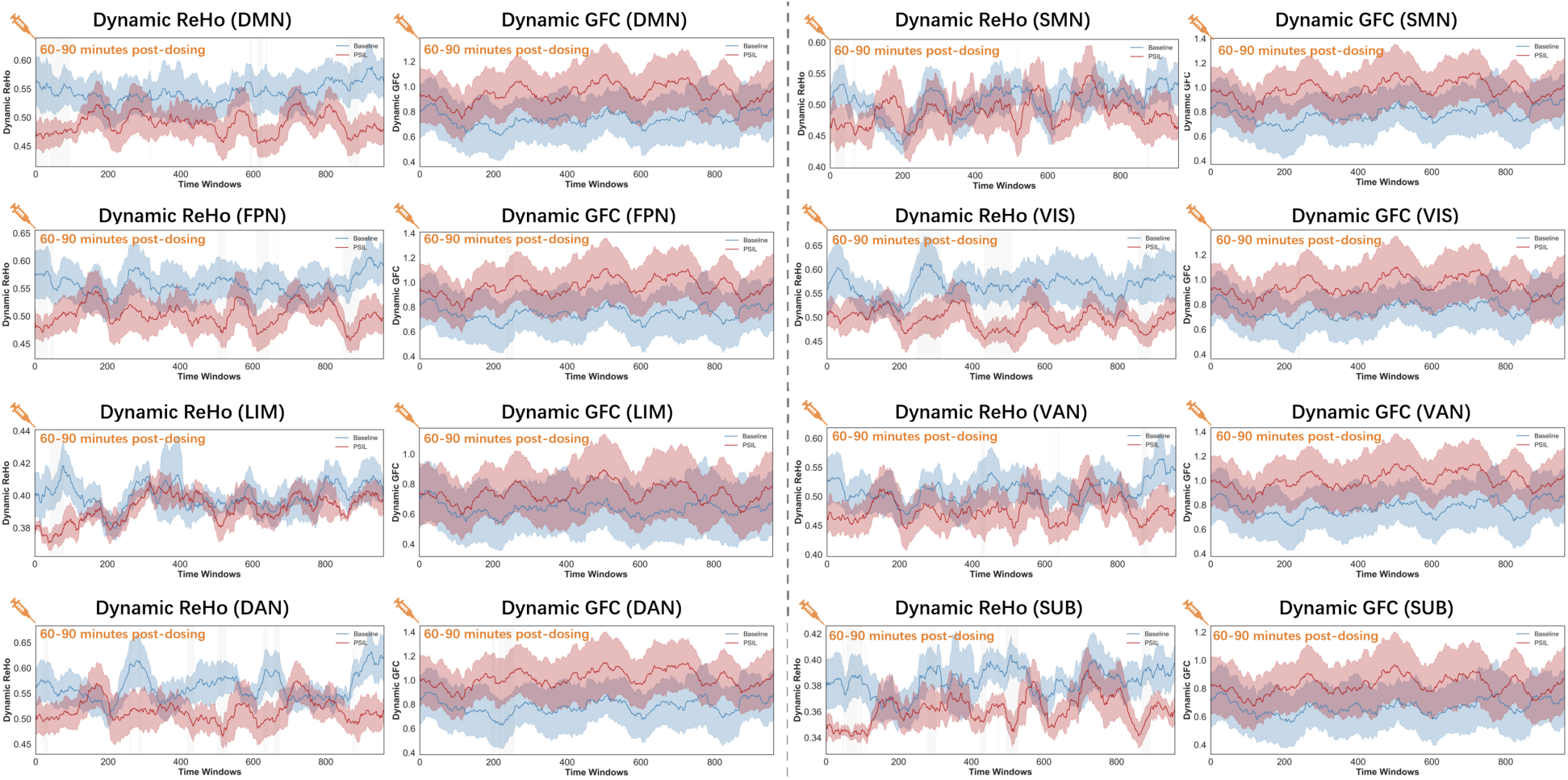
Dynamic regional homogeneity (ReHo) and global functional connectivity (GFC) in the psilocybin condition. Dynamic ReHo (left) and dynamic GFC (right) were computed using a sliding-window approach (window = 60 TRs, step = 1 TR) across the resting-state fMRI time series. Solid lines represent group mean values, with shaded bands indicating the standard error. Gray bars beneath each time course denote consecutive time windows where drug–baseline differences reached significance, as determined by cluster-based permutation testing (5,000 iterations).

**Figure S7:**
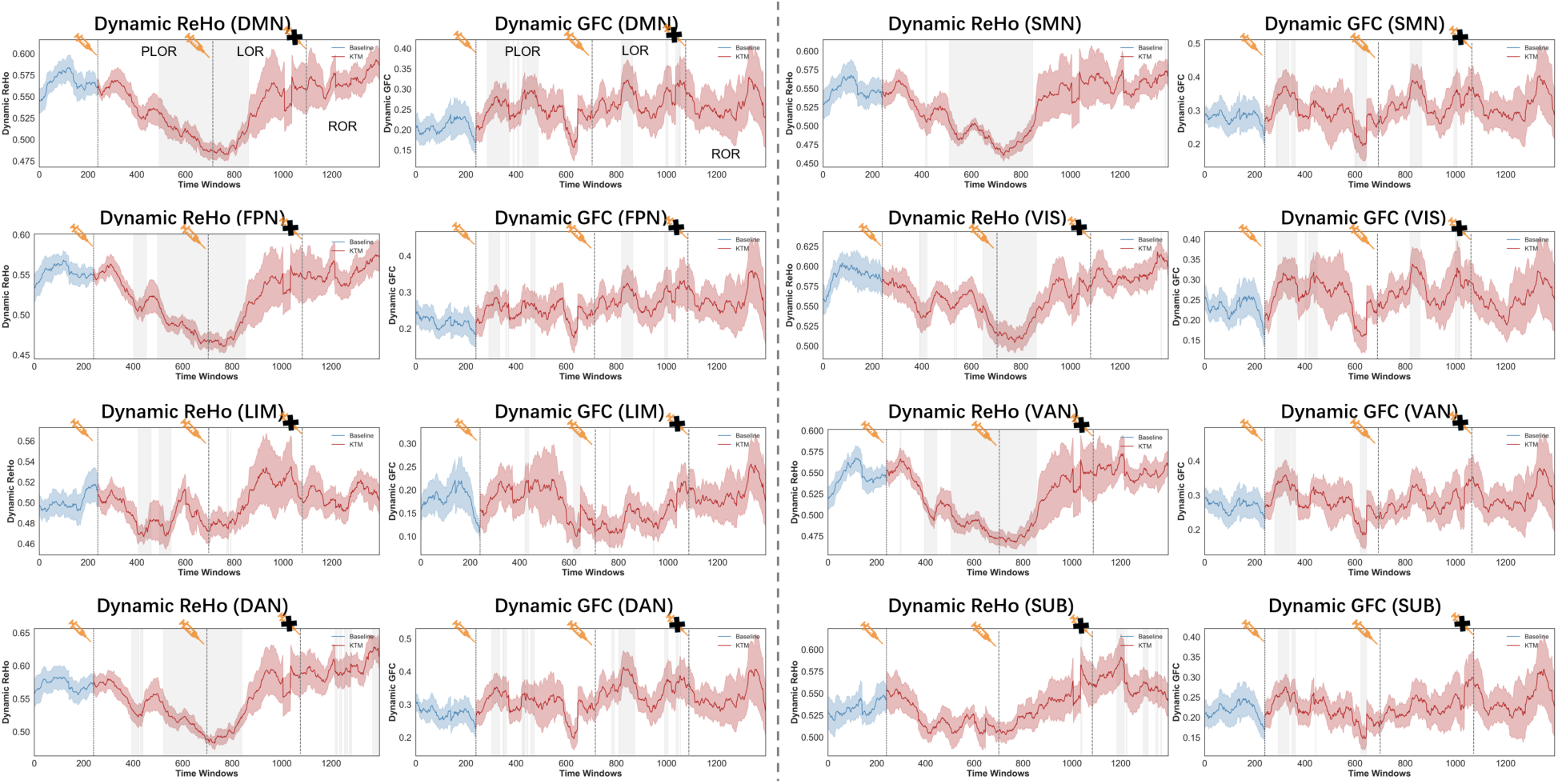
Dynamic regional homogeneity (ReHo) and global functional connectivity (GFC) in the ketamine condition. Dynamic ReHo (left) and dynamic GFC (right) were computed using a sliding-window approach (window = 60 TRs, step = 1 TR) across the resting-state fMRI time series. Solid lines represent group mean values, with shaded bands indicating the standard error. Gray bars beneath each time course denote consecutive time windows where drug–baseline differences reached significance, as determined by cluster-based permutation testing (5,000 iterations). PLOR = Pre-LOR (pre–loss of responsiveness), LOR = Loss of Responsiveness, ROR = Return of Responsiveness.

**Figure S8:**
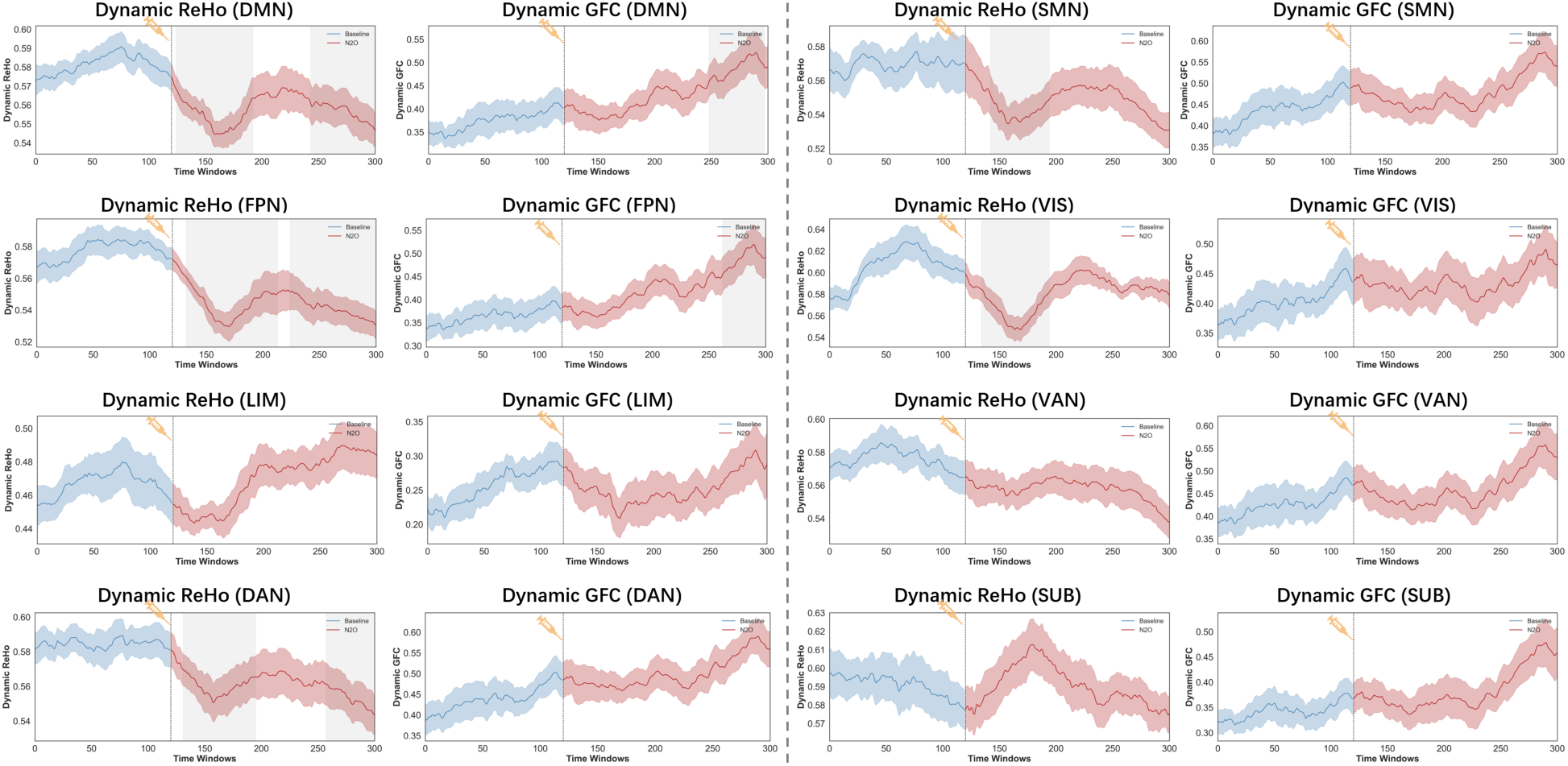
Dynamic regional homogeneity (ReHo) and global functional connectivity (GFC) in the nitrous oxide condition. Dynamic ReHo (left) and dynamic GFC (right) were computed using a sliding-window approach (window = 60 TRs, step = 1 TR) across the resting-state fMRI time series. Solid lines represent group mean values, with shaded bands indicating the standard error. Gray bars beneath each time course denote consecutive time windows where drug–baseline differences reached significance, as determined by cluster-based permutation testing (5,000 iterations).

**Figure S9:**
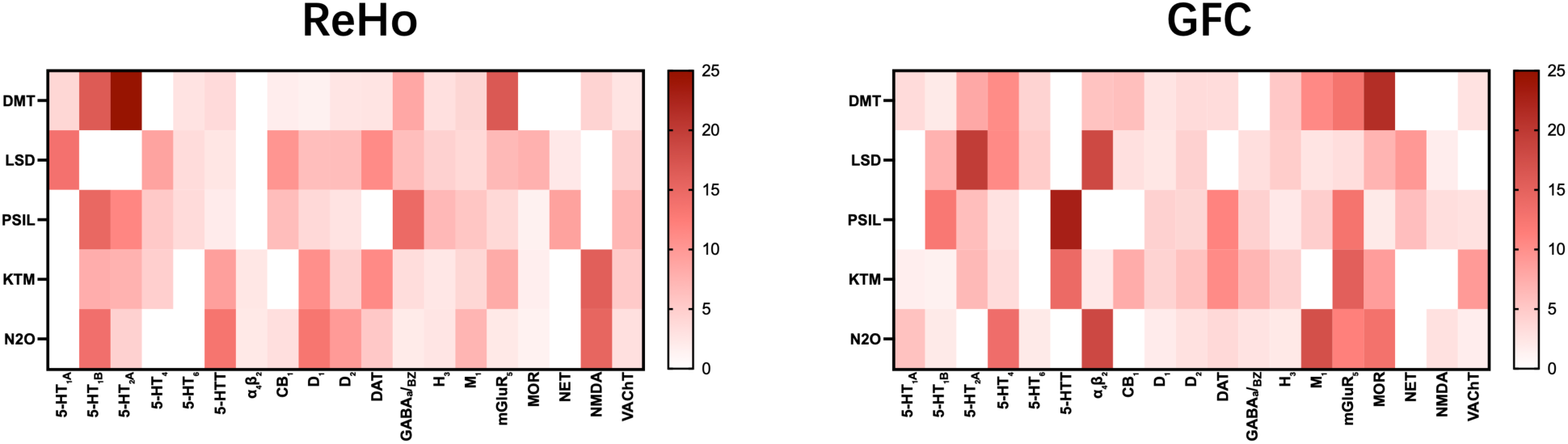
Comparative dominance analysis of neurotransmitter receptor contributions to regional homogeneity (ReHo) and global functional connectivity (GFC). Heatmaps display the relative contributions of 19 neurotransmitter receptors and transporter density maps in predicting psychedelic-induced brain activity changes. The left panel shows ReHo, and the right panel shows GFC. Rows indicate the five compounds (dimethyltryptamine/DMT, lysergic acid diethylamide/LSD, psilocybin/PSIL, ketamine/KTM, and nitrous oxide/N_2_O), and columns correspond to receptor or transporter systems. Color intensity represents the percentage contribution of each predictor to explained variance, as determined by dominance analysis.

**Figure S10:**
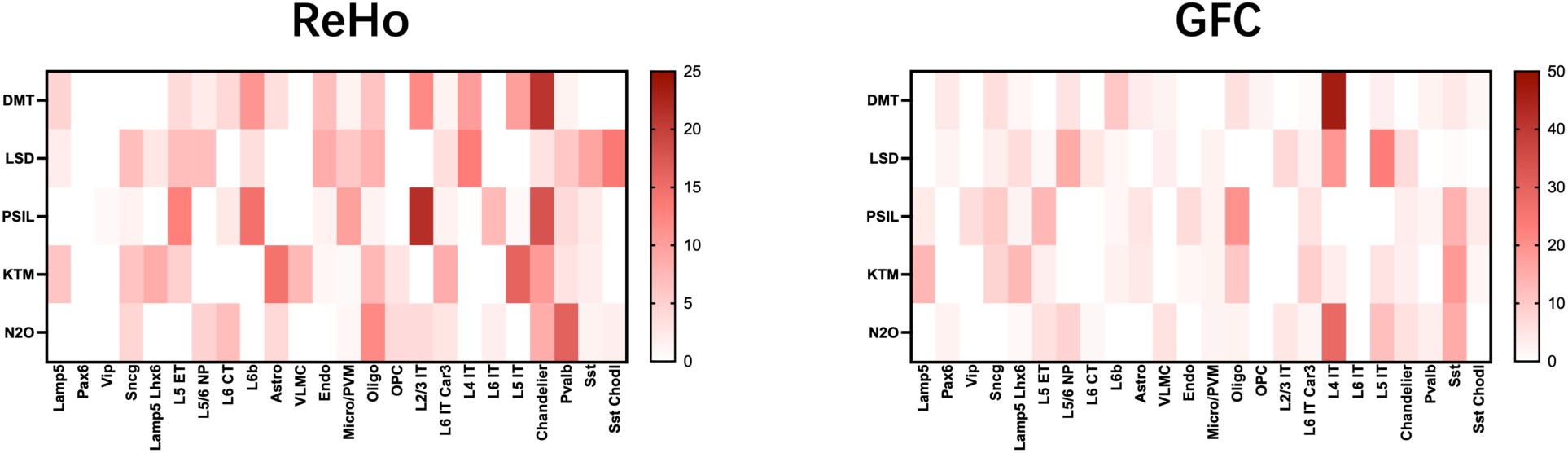
Comparative dominance analysis of cell type contributions to regional homogeneity (ReHo) and global functional connectivity (GFC). Heatmaps display the relative contributions of 24 cell type maps in predicting psychedelic-induced brain activity changes. The left panel shows ReHo, and the right panel shows GFC. Rows indicate the five compounds (dimethyltryptamine/DMT, lysergic acid diethylamide/LSD, psilocybin/PSIL, ketamine/KTM, and nitrous oxide/N_2_O), and columns correspond to cell types. Color intensity represents the percentage contribution of each predictor to explained variance, as determined by dominance analysis.

**Table S1.** DMT ReHo effect sizes and pFDR values across voxel neighborhood sizes.

| DMT | Global | SMN | VIS | DAN | VAN | LIM | FPN | DMN |
| --- | --- | --- | --- | --- | --- | --- | --- | --- |
| **7 vox** | d = 1.795,<br>pFDR < 0.001 | d = 0.952,<br>pFDR = 0.005 | d = 0.838,<br>pFDR = 0.010 | d = 1.570,<br>pFDR < 0.001 | d = 1.658,<br>pFDR < 0.001 | d = 0.142,<br>pFDR = 0.605 | d = 2.066,<br>pFDR < 0.001 | d = 2.278,<br>pFDR < 0.001 |
| **19 vox** | d = 1.820,<br>pFDR < 0.001 | d = 0.872,<br>pFDR = 0.009 | d = 0.875,<br>pFDR = 0.009 | d = 1.525,<br>pFDR < 0.001 | d = 1.562,<br>pFDR < 0.001 | d = 0.199,<br>pFDR = 0.469 | d = 2.029,<br>pFDR < 0.001 | d = 2.275,<br>pFDR < 0.001 |
| **27 vox** | d = 1.818,<br>pFDR < 0.001 | d = 0.863,<br>pFDR = 0.008 | d = 0.890,<br>pFDR = 0.007 | d = 1.493,<br>pFDR < 0.001 | d = 1.523,<br>pFDR < 0.001 | d = 0.224,<br>pFDR = 0.417 | d = 1.996,<br>pFDR < 0.001 | d = 2.241,<br>pFDR < 0.001 |
| **125 vox** | d = 1.548,<br>pFDR < 0.001 | d = 0.031,<br>pFDR = 0.911 | d = 0.897,<br>pFDR = 0.009 | d = 0.315,<br>pFDR = 0.260 | d = 1.215,<br>pFDR < 0.001 | d = 0.324,<br>pFDR = 0.247 | d = 1.837,<br>pFDR < 0.001 | d = 1.928,<br>pFDR < 0.001 |
| **343 vox** | d = 1.118,<br>pFDR = 0.001 | d = -0.336,<br>pFDR = 0.231 | d = 0.842,<br>pFDR = 0.008 | d = -0.125,<br>pFDR = 0.648 | d = 0.965,<br>pFDR = 0.003 | d = 0.401,<br>pFDR = 0.158 | d = 1.652,<br>pFDR < 0.001 | d = 1.704,<br>pFDR < 0.001 |
| **729 vox** | d = 0.730,<br>pFDR = 0.017 | d = -0.445,<br>pFDR = 0.120 | d = 0.767,<br>pFDR = 0.013 | d = -0.272,<br>pFDR = 0.328 | d = 0.703,<br>pFDR = 0.021 | d = 0.501,<br>pFDR = 0.083 | d = 1.476,<br>pFDR < 0.001 | d = 1.580,<br>pFDR < 0.001 |
| **1331 vox** | d = 0.474,<br>pFDR = 0.100 | d = -0.464,<br>pFDR = 0.106 | d = 0.695,<br>pFDR = 0.022 | d = -0.340,<br>pFDR = 0.226 | d = 0.431,<br>pFDR = 0.131 | d = 0.626,<br>pFDR = 0.036 | d = 1.288,<br>pFDR < 0.001 | d = 1.420,<br>pFDR < 0.001 |
| **2197 vox** | d = 0.326,<br>pFDR = 0.244 | d = -0.475,<br>pFDR = 0.099 | d = 0.637,<br>pFDR = 0.033 | d = -0.386,<br>pFDR = 0.173 | d = 0.215,<br>pFDR = 0.437 | d = 0.738,<br>pFDR = 0.016 | d = 1.113,<br>pFDR = 0.001 | d = 1.251,<br>pFDR < 0.001 |
| **3375 vox** | d = 0.232,<br>pFDR = 0.402 | d = -0.500,<br>pFDR = 0.084 | d = 0.593,<br>pFDR = 0.045 | d = -0.426,<br>pFDR = 0.135 | d = 0.064,<br>pFDR = 0.815 | d = 0.800,<br>pFDR = 0.010 | d = 0.970,<br>pFDR = 0.003 | d = 1.097,<br>pFDR = 0.001 |

**Table S2.** LSD ReHo effect sizes and pFDR values across voxel neighborhood sizes.

| LSD | Global | SMN | VIS | DAN | VAN | LIM | FPN | DMN |
| --- | --- | --- | --- | --- | --- | --- | --- | --- |
| **7 vox** | d = 2.137,<br>pFDR < 0.001 | d = 1.635,<br>pFDR < 0.001 | d = 2.243,<br>pFDR < 0.001 | d = 1.503,<br>pFDR < 0.001 | d = 1.153,<br>pFDR < 0.001 | d = 0.955,<br>pFDR = 0.002 | d = 2.275,<br>pFDR < 0.001 | d = 1.479,<br>pFDR < 0.001 |
| **19 vox** | d = 2.136,<br>pFDR < 0.001 | d = 1.619,<br>pFDR < 0.001 | d = 2.229,<br>pFDR < 0.001 | d = 1.438,<br>pFDR < 0.001 | d = 1.119,<br>pFDR < 0.001 | d = 1.134,<br>pFDR < 0.001 | d = 2.227,<br>pFDR < 0.001 | d = 1.455,<br>pFDR < 0.001 |
| **27 vox** | d = 2.131,<br>pFDR < 0.001 | d = 1.605,<br>pFDR < 0.001 | d = 2.227,<br>pFDR < 0.001 | d = 1.402,<br>pFDR < 0.001 | d = 1.101,<br>pFDR < 0.001 | d = 1.126,<br>pFDR < 0.001 | d = 2.197,<br>pFDR < 0.001 | d = 1.443,<br>pFDR < 0.001 |
| **125 vox** | d = 2.068,<br>pFDR < 0.001 | d = 1.476,<br>pFDR < 0.001 | d = 2.147,<br>pFDR < 0.001 | d = 1.112,<br>pFDR < 0.001 | d = 0.981,<br>pFDR = 0.002 | d = 0.911,<br>pFDR = 0.003 | d = 1.957,<br>pFDR < 0.001 | d = 1.349,<br>pFDR < 0.001 |
| **343 vox** | d = 1.901,<br>pFDR < 0.001 | d = 1.334,<br>pFDR < 0.001 | d = 1.922,<br>pFDR < 0.001 | d = 0.798,<br>pFDR = 0.009 | d = 0.899,<br>pFDR = 0.005 | d = 0.776,<br>pFDR = 0.009 | d = 1.650,<br>pFDR < 0.001 | d = 1.276,<br>pFDR < 0.001 |
| **729 vox** | d = 1.737,<br>pFDR < 0.001 | d = 1.213,<br>pFDR < 0.001 | d = 1.632,<br>pFDR < 0.001 | d = 0.481,<br>pFDR = 0.083 | d = 0.869,<br>pFDR = 0.006 | d = 0.789,<br>pFDR = 0.010 | d = 1.353,<br>pFDR < 0.001 | d = 1.201,<br>pFDR < 0.001 |
| **1331 vox** | d = 1.580,<br>pFDR < 0.001 | d = 1.112,<br>pFDR = 0.001 | d = 1.369,<br>pFDR < 0.001 | d = 0.196,<br>pFDR = 0.459 | d = 0.858,<br>pFDR = 0.006 | d = 0.874,<br>pFDR = 0.006 | d = 1.094,<br>pFDR = 0.001 | d = 1.091,<br>pFDR = 0.001 |
| **2197 vox** | d = 1.412,<br>pFDR < 0.001 | d = 1.017,<br>pFDR = 0.004 | d = 1.172,<br>pFDR = 0.002 | d = -0.025,<br>pFDR = 0.925 | d = 0.829,<br>pFDR = 0.007 | d = 0.954,<br>pFDR = 0.004 | d = 0.867,<br>pFDR = 0.006 | d = 0.962,<br>pFDR = 0.004 |
| **3375 vox** | d = 1.235,<br>pFDR = 0.002 | d = 0.921,<br>pFDR = 0.006 | d = 1.029,<br>pFDR = 0.004 | d = -0.179,<br>pFDR = 0.501 | d = 0.776,<br>pFDR = 0.013 | d = 1.020,<br>pFDR = 0.004 | d = 0.667,<br>pFDR = 0.025 | d = 0.852,<br>pFDR = 0.008 |

**Table S3.**
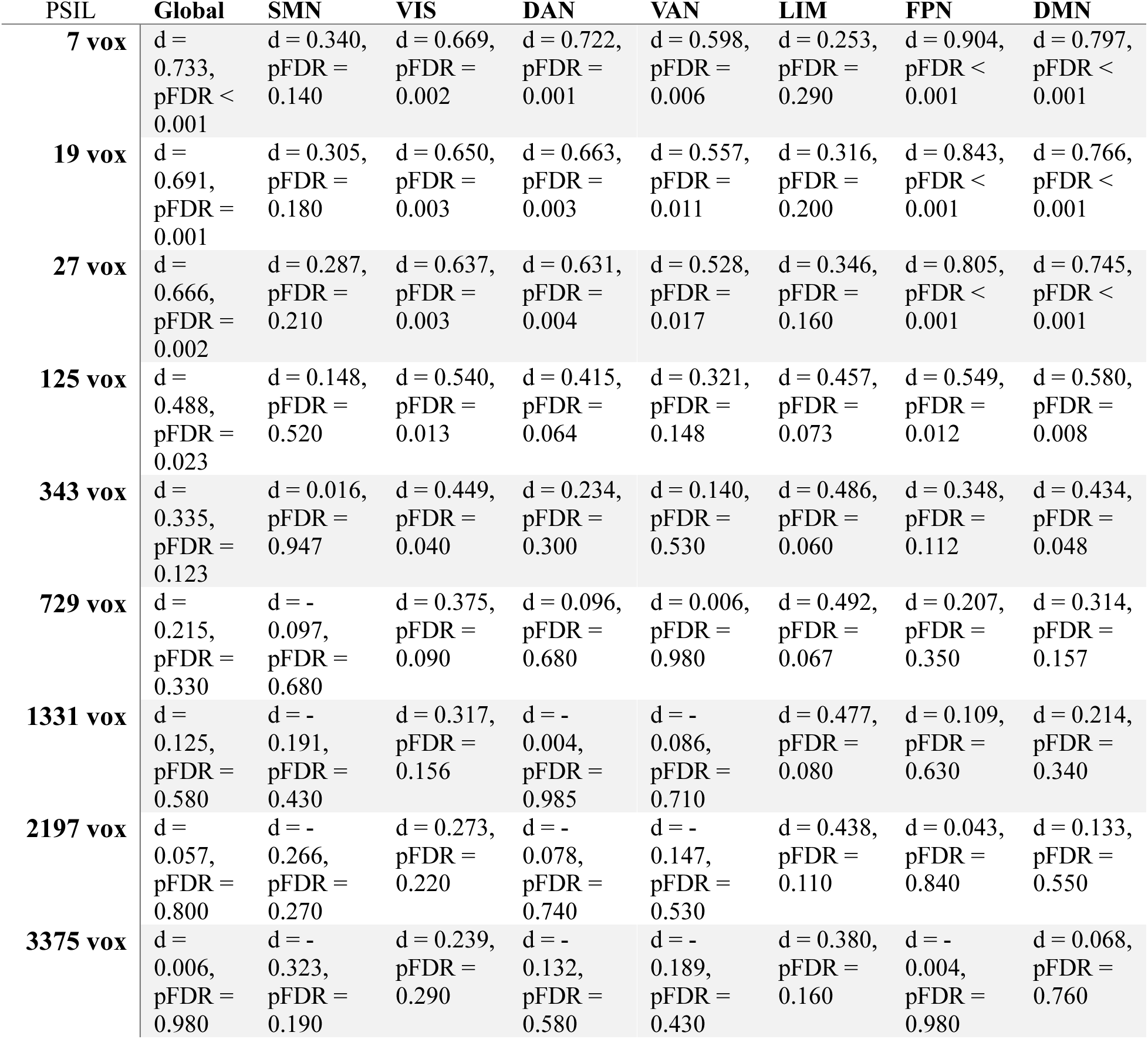
Psilocybin ReHo effect sizes and pFDR values across voxel neighborhood sizes.

**Table S4.** Ketamine ReHo effect sizes and pFDR values across voxel neighborhood sizes.

| KTM | Global | SMN | VIS | DAN | VAN | LIM | FPN | DMN |
| --- | --- | --- | --- | --- | --- | --- | --- | --- |
| **7 vox** | d = 0.733,<br>pFDR < 0.001 | d = 0.340,<br>pFDR = 0.140 | d = 0.669,<br>pFDR = 0.002 | d = 0.722,<br>pFDR = 0.001 | d = 0.598,<br>pFDR = 0.006 | d = 0.253,<br>pFDR = 0.290 | d = 0.904,<br>pFDR < 0.001 | d = 0.797,<br>pFDR < 0.001 |
| **19 vox** | d = 0.691,<br>pFDR = 0.001 | d = 0.305,<br>pFDR = 0.180 | d = 0.650,<br>pFDR = 0.003 | d = 0.663,<br>pFDR = 0.003 | d = 0.557,<br>pFDR = 0.011 | d = 0.316,<br>pFDR = 0.200 | d = 0.843,<br>pFDR < 0.001 | d = 0.766,<br>pFDR < 0.001 |
| **27 vox** | d = 0.666,<br>pFDR = 0.002 | d = 0.287,<br>pFDR = 0.210 | d = 0.637,<br>pFDR = 0.003 | d = 0.631,<br>pFDR = 0.004 | d = 0.528,<br>pFDR = 0.017 | d = 0.346,<br>pFDR = 0.160 | d = 0.805,<br>pFDR < 0.001 | d = 0.745,<br>pFDR < 0.001 |
| **125 vox** | d = 0.488,<br>pFDR = 0.023 | d = 0.148,<br>pFDR = 0.520 | d = 0.540,<br>pFDR = 0.013 | d = 0.415,<br>pFDR = 0.064 | d = 0.321,<br>pFDR = 0.148 | d = 0.457,<br>pFDR = 0.073 | d = 0.549,<br>pFDR = 0.012 | d = 0.580,<br>pFDR = 0.008 |
| **343 vox** | d = 0.335,<br>pFDR = 0.123 | d = 0.016,<br>pFDR = 0.947 | d = 0.449,<br>pFDR = 0.040 | d = 0.234,<br>pFDR = 0.300 | d = 0.140,<br>pFDR = 0.530 | d = 0.486,<br>pFDR = 0.060 | d = 0.348,<br>pFDR = 0.112 | d = 0.434,<br>pFDR = 0.048 |
| **729 vox** | d = 0.215,<br>pFDR = 0.330 | d = -0.097,<br>pFDR = 0.680 | d = 0.375,<br>pFDR = 0.090 | d = 0.096,<br>pFDR = 0.680 | d = 0.006,<br>pFDR = 0.980 | d = 0.492,<br>pFDR = 0.067 | d = 0.207,<br>pFDR = 0.350 | d = 0.314,<br>pFDR = 0.157 |
| **1331 vox** | d = 0.125,<br>pFDR = 0.580 | d = -0.191,<br>pFDR = 0.430 | d = 0.317,<br>pFDR = 0.156 | d = -0.004,<br>pFDR = 0.985 | d = -0.086,<br>pFDR = 0.710 | d = 0.477,<br>pFDR = 0.080 | d = 0.109,<br>pFDR = 0.630 | d = 0.214,<br>pFDR = 0.340 |
| **2197 vox** | d = 0.057,<br>pFDR = 0.800 | d = -0.266,<br>pFDR = 0.270 | d = 0.273,<br>pFDR = 0.220 | d = -0.078,<br>pFDR = 0.740 | d = -0.147,<br>pFDR = 0.530 | d = 0.438,<br>pFDR = 0.110 | d = 0.043,<br>pFDR = 0.840 | d = 0.133,<br>pFDR = 0.550 |
| **3375 vox** | d = 0.006,<br>pFDR = 0.980 | d = -0.323,<br>pFDR = 0.190 | d = 0.239,<br>pFDR = 0.290 | d = -0.132,<br>pFDR = 0.580 | d = -0.189,<br>pFDR = 0.430 | d = 0.380,<br>pFDR = 0.160 | d = -0.004,<br>pFDR = 0.980 | d = 0.068,<br>pFDR = 0.760 |

**Table S5.** Nitrous oxide ReHo effect sizes and pFDR values across voxel neighborhood sizes.

| N<sub>2</sub>O | Global | SMN | VIS | DAN | VAN | LIM | FPN | DMN |
| --- | --- | --- | --- | --- | --- | --- | --- | --- |
| **7 vox** | d = 1.304,<br>pFDR < 0.001 | d = 0.871,<br>pFDR = 0.005 | d = 1.051,<br>pFDR = 0.001 | d = 1.225,<br>pFDR < 0.001 | d = 0.922,<br>pFDR = 0.003 | d = -0.366,<br>pFDR = 0.178 | d = 1.536,<br>pFDR < 0.001 | d = 1.084,<br>pFDR = 0.001 |
| **19 vox** | d = 1.205,<br>pFDR < 0.001 | d = 0.821,<br>pFDR = 0.007 | d = 0.973,<br>pFDR = 0.002 | d = 1.105,<br>pFDR < 0.001 | d = 0.816,<br>pFDR = 0.007 | d = -0.359,<br>pFDR = 0.186 | d = 1.445,<br>pFDR < 0.001 | d = 1.037,<br>pFDR = 0.001 |
| **27 vox** | d = 1.143,<br>pFDR < 0.001 | d = 0.791,<br>pFDR = 0.008 | d = 0.920,<br>pFDR = 0.003 | d = 1.038,<br>pFDR = 0.001 | d = 0.750,<br>pFDR = 0.012 | d = -0.360,<br>pFDR = 0.185 | d = 1.383,<br>pFDR < 0.001 | d = 1.004,<br>pFDR = 0.002 |
| **125 vox** | d = 0.677,<br>pFDR = 0.020 | d = 0.580,<br>pFDR = 0.041 | d = 0.484,<br>pFDR = 0.082 | d = 0.555,<br>pFDR = 0.050 | d = 0.308,<br>pFDR = 0.252 | d = -0.355,<br>pFDR = 0.191 | d = 0.945,<br>pFDR = 0.003 | d = 0.700,<br>pFDR = 0.017 |
| **343 vox** | d = 0.236,<br>pFDR = 0.375 | d = 0.356,<br>pFDR = 0.190 | d = 0.072,<br>pFDR = 0.786 | d = 0.131,<br>pFDR = 0.619 | d = -0.023,<br>pFDR = 0.930 | d = -0.336,<br>pFDR = 0.214 | d = 0.541,<br>pFDR = 0.055 | d = 0.314,<br>pFDR = 0.244 |
| **729 vox** | d = -0.071,<br>pFDR = 0.786 | d = 0.151,<br>pFDR = 0.569 | d = -0.181,<br>pFDR = 0.495 | d = -0.144,<br>pFDR = 0.587 | d = -0.238,<br>pFDR = 0.372 | d = -0.330,<br>pFDR = 0.222 | d = 0.201,<br>pFDR = 0.450 | d = -0.015,<br>pFDR = 0.954 |
| **1331 vox** | d = -0.258,<br>pFDR = 0.335 | d = -0.019,<br>pFDR = 0.943 | d = -0.309,<br>pFDR = 0.252 | d = -0.303,<br>pFDR = 0.261 | d = -0.371,<br>pFDR = 0.173 | d = -0.329,<br>pFDR = 0.224 | d = -0.058,<br>pFDR = 0.826 | d = -0.229,<br>pFDR = 0.390 |
| **2197 vox** | d = -0.370,<br>pFDR = 0.174 | d = -0.153,<br>pFDR = 0.563 | d = -0.374,<br>pFDR = 0.169 | d = -0.390,<br>pFDR = 0.153 | d = -0.457,<br>pFDR = 0.098 | d = -0.330,<br>pFDR = 0.222 | d = -0.242,<br>pFDR = 0.364 | d = -0.359,<br>pFDR = 0.186 |
| **3375 vox** | d = -0.439,<br>pFDR = 0.111 | d = -0.256,<br>pFDR = 0.338 | d = -0.415,<br>pFDR = 0.131 | d = -0.436,<br>pFDR = 0.113 | d = -0.512,<br>pFDR = 0.067 | d = -0.334,<br>pFDR = 0.217 | d = -0.365,<br>pFDR = 0.179 | d = -0.443,<br>pFDR = 0.108 |

